# Developmental Timing Establishes Hematopoietic Stem and Progenitor Cell Lineage Bias

**DOI:** 10.1101/2025.05.12.653589

**Authors:** Anastasia Nizhnik, Bianca A. Ulloa, Kevyn Jackson, Deyou Zheng, Teresa V. Bowman

## Abstract

Embryonic development is a critical time window for the establishment of the hematopoietic and immune systems. During embryogenesis, hematopoietic stem and progenitor cells (HSPCs) that possess divergent differentiation preferences arise and sustain lifelong hematopoiesis. How the pace and differentiation repertoire of native hematopoiesis is set in development and how it evolves as the organism grows is incompletely elucidated. Here, we use temporal lineage tracing of the emerging hematopoietic system during zebrafish development coupled with single cell RNA sequencing to define the origins of larval and adult hematopoietic cells. In both larvae and adult tissues, HSPCs arising earlier during embryogenesis show a predilection for lymphoid lineage production while those arising later are skewed towards erythroid. Moreover, early arising HSPCs are the main contributors to adult tissue resident lymphocytes. Mechanistically, early and late arising HSPCs show divergent transcriptional signatures and differential sensitivities to the levels of the transcription factor Runx1. Additionally, we identified that young larvae possess a more heterogeneous set of lymphoid cells than previously recognized including diverse T-lymphocytes and Innate Lymphoid-like Cells (ILCs). We demonstrated that these newly described larval ILC-like cells reside in lymphoid and mucosal organs and are responsive to viral mimic immune stimulation, indicating their relevance in early vertebrate life. The work provides new fundamental knowledge on how the heterogeneous HSPC pool establishes early immune hierarchies during embryogenesis and its persistence in adulthood.

## INTRODUCTION

The hematopoietic system is comprised of an array of blood cells carrying out a variety of functions ranging from oxygenation to pathogen defense. These differentiated mature blood cells are produced by Hematopoietic Stem and Progenitor Cells (HSPCs) that possess multilineage differentiation potential. Although multipotency is a defining feature of HSPCs, their differentiation output is not always balanced. HSPCs can have lineage preferences or biases in the type of mature cells they make with observed predilections toward lymphoid, myeloid, or erythroid lineages.^1–3^ HSPC lineage preferences are evident and important in human health and seem to have an age dependency to their features. For example, age-dependent myeloid skewing seems to be attributed in part to shifts in the relative frequency and function of lymphoid and myeloid biased HSPC.^4^ Additionally, lymphoid leukemias are more common in childhood, while myeloid leukemias are more common with age. Although the mechanisms guiding lineage preference of HSPCs remains elusive, recent work suggests that it could originate embryonically and persist throughout life.^5^

Embryonic development is a critical stage for the formation of the hematopoietic system, where multiple sequential and overlapping ‘waves’ of hematopoiesis produce distinct HSPCs and mature cell types.^6–8^ The cells generated at each wave serve a particular purpose for the proper establishment and function of lifelong hematopoietic and immune systems.^9^ Currently, the field refers to two main waves of hematopoiesis: primitive and definitive.^8^ The ‘primitive wave’ gives rise to macrophages, erythrocytes, and neutrophils.^10,11^ The ‘definitive wave’ is subdivided into an earlier pro-definitive phase and a definitive phase.^9^ The pro-definitive phase progenitors generate myeloid lineage cells, megakaryocytes, B-cells, and NK cells.^9,12–14^ The definitive phase produces Hematopoietic Stem Cells (HSCs) that can sustain lifelong hematopoiesis. The classic paradigm was that pro-definitive wave cells were transient and only necessary for fetal life, while definitive wave HSCs sustained the entirety of the adult hematopoietic and immune systems. Work over the past decade using lineage tracing methods that permit dissection of native hematopoiesis have challenged this model and illustrated that pro-definitive wave progenitors can contribute to adult steady-state bone marrow hematopoiesis,^5^ tissue-resident macrophages,^15,16^ and innate B1a cells.^17^ These paradigm-challenging studies illuminated that much remains unknown about the differentiation repertoire of the heterogeneous pool of embryonically emerging HSPCs and the persistence of their progeny beyond development.

While the embryonic origins of tissue-resident macrophages and B-1a lymphocytes are studied,^7,18^ hematopoietic origins of other Innate Lymphoid Cells (ILCs) are poorly resolved. ILCs belong to the lymphoid lineage and resemble T-helper lymphocytes, however ILCs lack the adaptive antigen-specific receptors characteristic of T lymphocytes.^19^ ILCs are grouped into three categories: ILC1, ILC2, and ILC3, paralleling the T-helper groups Th1, Th2, and Th17, respectively.^20^ In adult tissues, ILCs can secrete cytokines and recruit other immune cells in response to infections but their behaviors in perinatal life is poorly resolved.^21^ Moreover, it is unknown currently if the lineage biases of embryonic HSPCs is relevant to ILC ontogeny.

While substantial work in the field dissected the differences between extraembryonic-derived HSC-independent progenitors and intraembryonic-derived HSCs, there is less research on the influence of developmental timings on HSPC properties. To address this gap, we sought to resolve the cell types generated by early *versus* late emerging HSPCs during zebrafish development using a temporally inducible lineage tracing system followed by single cell RNA sequencing (scRNA-seq). By lineage-tracing HSPC progeny in larval and adult life, we find that embryonic time imparts discrete and long-lasting differentiation functions onto HSPCs: early arising HSPCs produce more lymphoid lineage cells and later arising HSPCs show a preference towards the erythroid lineage. The lymphoid bias of early arising HSPCs in adult organs was evident not only in marrow but also in barrier tissues like the skin and intestine. Mechanistically, early- and late-emerging HSPCs exhibit differential sensitivity to the levels of the transcription factor Runx1. Furthermore, we uncovered that HSPCs make a diverse set of previously undescribed immune cells in larval zebrafish, including various groups of ILCs and several other novel immune cells that are responsive to immune stimulation in early life. Overall, the study presents new insights on the embryonic regulation of HSPC lineage bias across a broad swath of blood and immune tissues suggesting that embryonic environment could have long lasting impacts on hematopoietic aging and chronic inflammatory disease.

## RESULTS

### Lineage tracing and scRNA-seq resolve early, mid and late HSPCs in zebrafish

During embryonic and larval development, long-lived HSCs and HSC-independent progenitors display distinct lymphoid-myeloid differentiation kinetics.^22–24^ HSC-independent progenitors are the first to differentiate into mature immune cells, while HSC-derived progeny appear later.^22–24^ While prior studies established this new paradigm, they were limited in the cell types they could examine due to technical limitations. The hematopoietic system is diverse, with numerous immune cell types that serve an array of functions and are found localized in many tissues. In a previous zebrafish lineage tracing study^22^, neutrophils and thymic cells were examined, but it is unclear if the differentiation repertoire and kinetics that distinguish HSC and HSC-independent progenitors in larvae is unique to these cell types or more generalizable throughout the hematopoietic system. To address this question, we used an inducible Cre-Lox-based genetic lineage tracing approach that fluorescently labels HSPCs and their progeny, coupled with scRNA-seq to delineate the identities of labeled cells (**Fig. 1A**). The Cre-Lox system is comprised of two components: *Tg(draculin:creER^T^*^2^), which allows hematopoietic-enriched expression of a tamoxifen-inducible Cre recombinase, and *Tg(ubi:loxP-GFP-loxP-stop-mCherry),* which allows for ubiquitous expression of fluorescent proteins.^25,26^ Upon addition of tamoxifen, Cre recombinase excises the loxP-GFP-loxP cassette, allowing for the expression of mCherry. The result is permanent mCherry labeling of *draculin+* cells, including HSPCs and their differentiated progeny. From prior studies employing the *drl:CreER^T2^* system, it will label HSPCs contributing to larval hematopoiesis including early thymic-seeding cells^22^ as well as long-lived HSPCs that sustain all blood lineages within the adult kidney marrow (mammalian bone marrow equivalent),^27^ suggesting it encompasses both HSC-independent progenitor and HSC populations. We think this could include the previously described erythroid-myeloid progenitors based on expression of *drl-*driven transgenes within the microenvironment where these cells reside during development.^22^ Previously, we determined that varying the window of tamoxifen exposure during embryonic development helped to distinguish HSPC differentiation kinetics into thymic cells and neutrophils.^22^ In the current study, tamoxifen-mediated Cre induction of mCherry labelling is begun at either one, two, or three days post fertilization (dpf), with 4-hydroxytamoxifen added for 20 hours (**Fig. 1A**). We refer to the labeling at one dpf as “early-HSPCs”, at two dpf as “mid-HSPCs” and at three dpf as “late-HSPCs”. These time points were chosen as they represent early, middle, and late time points during HSPC specification, maturation, and expansion during zebrafish development.^6^ Based on prior studies,^22–24^ we hypothesized that early-HSPCs would be enriched for cells with rapid lymphoid differentiation properties and mid/late-HSPCs would be enriched for long-lived HSCs.^22^ To identify what cells are labeled at early, mid, and late induction time points, we performed scRNA-seq on mCherry-expressing cells isolated from larvae one-day-post labelling, so at two, three, and four dpf, respectively. To assess the subsequent differentiated hematopoietic progeny, mCherry-expressing cells were sequenced from 6 and 10 dpf larvae (**Fig. 1A**). After filtering out low quality cells based on standard quality control parameters in the field,^28^ we clustered cells in Seurat.^29^ In total, about 75% of mCherry-expressing cells were identified as endothelial or hematopoietic, while the remaining 25% were non-hematopoietic cells like mesenchyme, neurons, cardiac, muscle, and epidermis and thus were filtered out for subsequent analyses (**Supplementary Fig. 1A-D, Table S1**). As a result, 25 hematopoietic and immune clusters were detected across all timepoints and traces, including HSCs, progenitors, and other cells of the major mature hematopoietic lineages, including lymphoid, myeloid, and erythroid cells (**Fig. 1B, Supplementary Fig. 1E,F & 2A-C, Table S1**). To facilitate ease of access for the community, this full dataset and all others described below are available to search via a shinyapp webtool made with ShinyCell^30^ at https://sites.google.com/view/tvbowmanlab/developmental-hematopoiesis-atlas.

**Figure 1.**
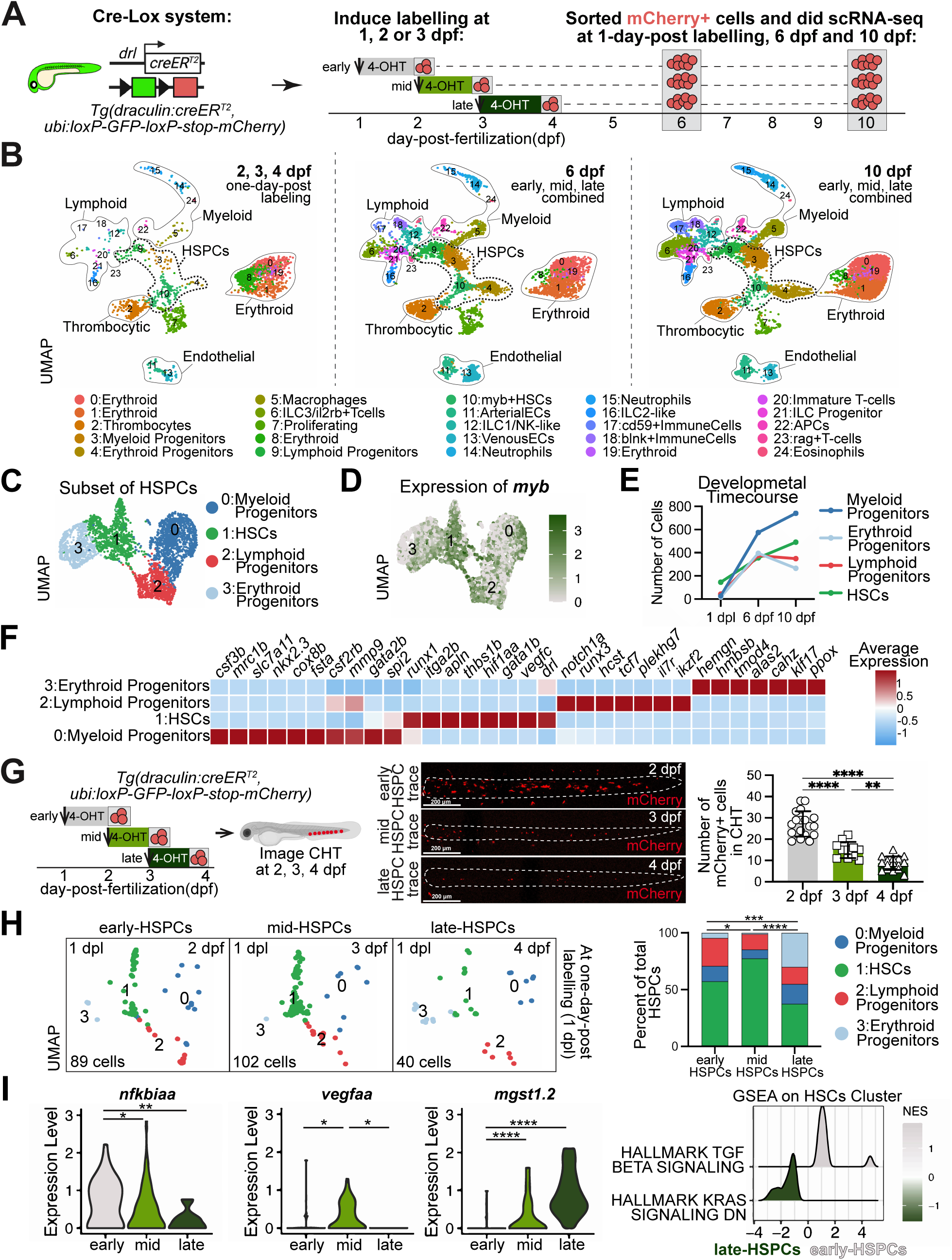
Lineage tracing of early, mid, and late HSPCs in zebrafish development. (**A**) Inducible Cre-Lox labelling system was activated with 12 𝜇M hydroxy-tamoxifen (4-OHT) at either 1, 2, or 3 days-post-fertilization (dpf), termed “early-HSPCs”, “mid-HSPCs” and “late-HSPCs,” respectively. scRNA-seq of mCherry-expressing progeny of each cohort/trace was done at three developmental timepoints: one-day-post labelling (1 dpl), 6 dpf, and 10 dpf. (**B**) UMAP showing hematopoietic clusters derived from early, mid, and late HSPCs across all tested time points. (**C**) UMAP showing the HSPC subsets in all traces. (**D**) Feature plots of *myb* expression across HSPC clusters. (**E**) Line plot showing the quantification of larval HSPCs detected by scRNA-seq in all traces. (**F**) Heatmap showing select genes in the HSPC subsets from a 4-way differential expression analysis. (**G**) Confocal imaging of mCherry signal in the CHT at one-day-post labeling for each tamoxifen induced labeling timepoint. Graph showing reduced mCherry signal at 3 & 4 dpf (mid & late trace) timepoints compared to the 2 dpf (early trace) signal. Each point represents one fish. Significance computed with one-way ANOVA with Tukey multiple testing corrections. (**H**) UMAP of 1 dpl HSPCs and their relative percentages in the bar plot. (**I**) Violin plots showing the trace-selective expression of genes in cluster 10 myb+HSCs. Significance computed with one-way ANOVA with Tukey multiple testing corrections. Plot on the left is GSEA analysis of the same cluster. *p<0.05, **p ≤0.01, **** p ≤0.0001

The HSPC clusters were identified based on the expression of *myb*, which is a marker gene commonly used to assess HSPCs by *in situ* hybridization.^31,32^ Expression of *myb* across all time points was localized to four stem and progenitor clusters identified as Myeloid Progenitors (MyPs), Erythroid Progenitors (EryPs), Lymphoid Progenitors (LyPs), and HSCs (**Fig. 1C,D**). The number of all HSPCs increased with developmental time (**Fig. 1E**). Differential expression analysis comparing HSCs to EryPs, MyPs and LyPs showed distinct transcriptomes across the four clusters (**Fig. 1F, Table S2**). Consistent with the lineage-enriched expression, EryPs express higher levels of erythroid genes such as *klf17, alas2*, and *hemgn*, MyPs express higher levels of genes critical for the myeloid lineage such as *spi2, csf2rb* and *mmp9*, and LyPs express higher levels of genes associated with lymphopoiesis such as *runx3, tcf7,* and *il7r* (Table S2). HSCs were enriched for genes encoding stem cell factors like *runx1*, *drl*, *apln,* and *itga2b*.

To identify transcriptional changes occurring in HSPCs as they mature during development, we did gene enrichment analysis comparing each cluster over developmental time (**Supplementary Fig. 2D, Table S2**). We observed that the HSC transcriptome was enriched for genes involved in RNA splicing and metabolism at 1-day-post labeling and 10 dpf, with elevated expression of genes involved with hypoxia at 6 dpf. In contrast, LyPs and MyPs showed more similar changes with elevated expression of genes involved with proteostasis, protein translation, and protein folding at 1-day-post labeling and 10 dpf. These observations are in line with the rapid growth and protein production typically associated with hematopoietic progenitors.^33^

To visualize the early-, mid-, and late-HSPCs *in vivo*, we did confocal imaging of the larval zebrafish HSPC niche termed the Caudal Hematopoietic Tissue (CHT) at 1-day-post labelling (**Fig. 1G**). The CHT contained the highest number of mCherry-labeled cells in the early-HSPC trace and the lowest number in the late-HSPC trace. To identify if there were differences in the composition of HSPCs labeled by each trace, we analyzed the HSPC subsets at 1-day-post labeling in the scRNA-seq dataset (**Fig. 1H**). Although all four populations, MyPs, LyPs, EryPs and HSCs, were detected in the early-, mid-, and late-traces, the frequency of the populations was distinct. LyPs were most abundant in the early-HSPC trace, HSCs were enriched in the mid-HSPC trace, while EryPs were most frequent in the late-HSPC trace. Comparison of differentially expressed genes within the *myb+* HSC cluster at 1-day-post labeling from early-, mid-, and late-HSPC traces uncovered differences related to lineage and signaling (**Fig. 1I**). These data highlight differences within the HSPC pools across the window of HSPC specification, maturation, and expansion during zebrafish development.

### HSPCs differentiate into a large repertoire of immune and hematopoietic cell types during larval life

By 6 and 10 dpf, HSPCs differentiate into various immune and hematopoietic lineages. Consistent with our prior study,^22^ there is a substantial contribution of the early-HSPCs to the CHT and thymus at 6 dpf, with minimal contributions from mid- and late-HSPC traces, as observed by confocal imaging (**Fig. 2A,B**). The precise identity of the hematopoietic cells arising in the hematopoietic tissues was unclear, so we leveraged the scRNA-seq data to identify the immune cells derived from early-, mid-, and late-HSPC lineage traces. Lymphoid and myeloid lineage clusters were subset and analyzed by developmental timepoint (**Fig. 2C, Table S3**). When examined as all lymphoid clusters in aggregate, we observed a burst of differentiation from the early-HSPCs, and a concomitant decrease in the proportion of LyPs, potentially due to ongoing differentiation (**Fig. 2D**). Compared to mid- and late-HSPCs, the early-HSPCs showed a stronger propensity for differentiation into lymphoid cluster 3 identified as ILC3/il2rb+Tcells. This observation parallels confocal imaging, where early-HSPCs contributed significantly more to the thymus than mid- or late-HSPCs (**Fig. 2A**). Not all lymphoid clusters behaved the same. For example, cluster 13 rag+ T cells arose at similar frequency from early- and mid-HSPCs, while late-HSPCs did not generate rag+ T cells by 10 dpf (**Fig. 2D**). In contrast to the burst of lymphoid production, analysis of all myeloid lineage clusters in aggregate showed a steady level of differentiation from all three HSPC traces (**Fig. 2E**). However, the production of myeloid lineage cells by the three HSPC pools was not entirely homogenous. Late-HSPCs showed a bias toward making more MyPs at 6 and 10 dpf compared to early- or mid-HSPCs, suggesting that HSPCs maybe skewed toward myeloid lineage differentiation during later larval development. Furthermore, macrophages and Antigen Presenting Cells (APCs) showed different enrichments among the three traces (**Fig. 2E**). In sum, the confocal imaging data along with single cell RNA sequencing illustrate that early HSPCs are preferentially differentiating to lymphocytes (**Fig. 2F**).

**Figure 2.**
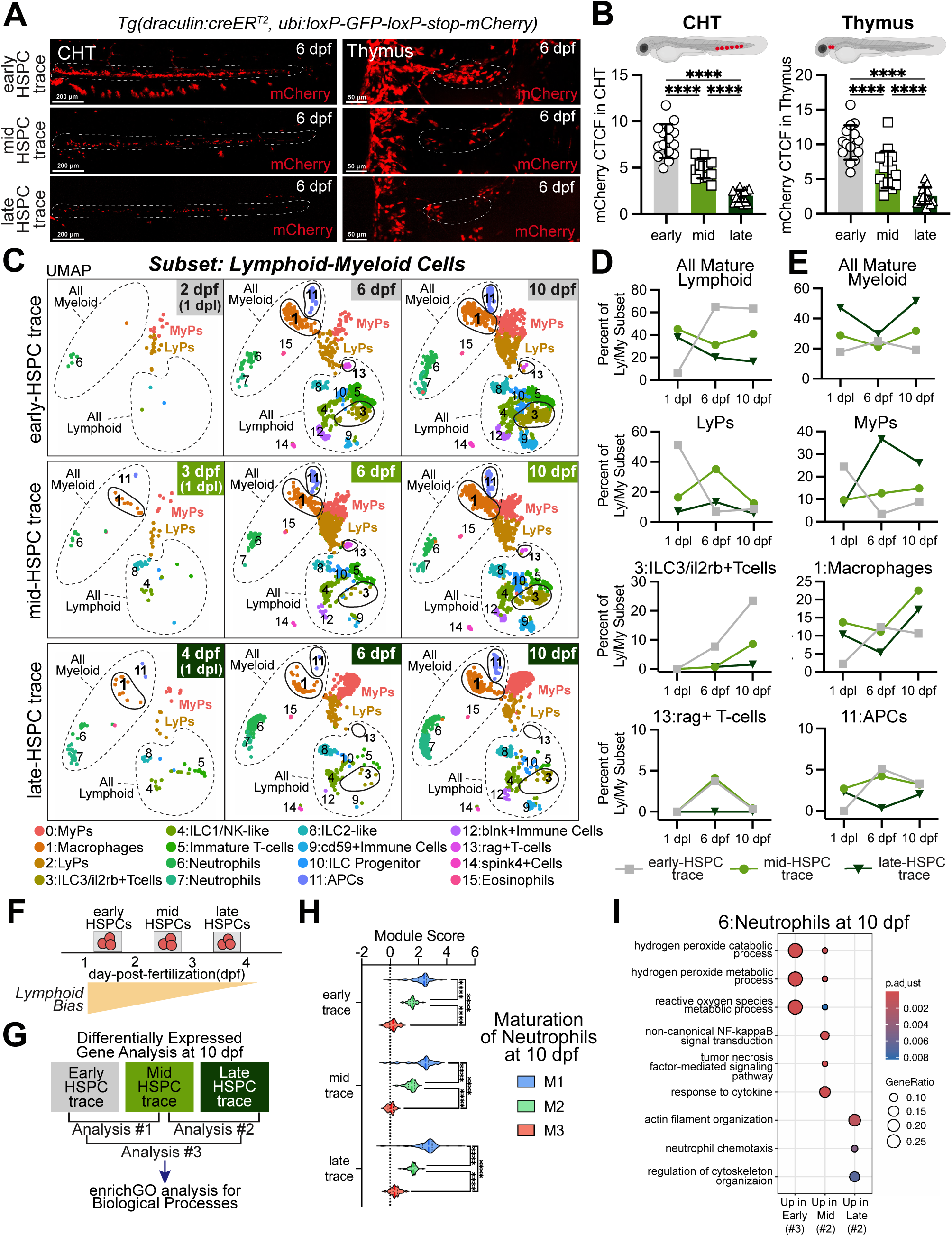
Differentiation of early, mid, and late HSPCs to myeloid and lymphoid lineages during development. (**A**) Confocal imaging of mCherry signal derived from early, mid, and late HSPCs at 6dpf with the thymus and caudal hematopoietic tissue (CHT) circled with dotted white lines. (**B**) Bar plots showing hither CHT and thymus signal from early HSPCs. Each point represents one fish. Significance computed with one-way ANOVA with Tukey multiple testing corrections. (**C**) UMAP showing the lymphoid-myeloid subset of cells, separated by timepoint and trace with dotted lines around the myeloid and lymphoid lineage cells and solid lines around clusters highlighted in D,E. (**D,E**) Plots show percentage of cells in a given cluster by trace across developmental timepoints. Lymphoid cells shown in (D) and Myeloid cells shown in (E). (**F**) Schema of the proposed model of lymphoid lineage bias established by developmental timing. (**G**) Experimental schema of differential gene expression between 10dpf cells derived from early, mid, and late HSPCs. (**H**) Module scoring of maturation stage (M1-M3) from least to most mature neutrophils across the early, mid, and late traces. Significance computed with one-way ANOVA with Tukey multiple testing corrections. (**I**) Biological Process EnrichGO pathway enrichment analysis showing pathways enriched in 10 dpf Neutrophils. **** p ≤0.0001

Immune cells derived from distinct developmental sources can have transcriptomes that reflect different functions.^34^ To determine if the immune cells derived from early-, mid- and late-HSPCs are transcriptionally distinct, we did differential gene expression and pathway enrichment analyses (**Fig. 2G**). The analysis was focused on neutrophils because there are established gene signatures in zebrafish that can allow us to assess if transcriptional differences are due to neutrophil maturation stage or reflect potential functional differences related to the originating HSPC. First, we examined maturation stage based on M1 (immature) to M3 (most mature) gene module scores defined by Kirchberger et al.^35^ We compared the neutrophils from all three traces at 10dpf as we posited these would be at similar maturation stages. The module score analysis confirmed that hypothesis with no major differences in the distribution of neutrophil maturation stages across the three traces with the majority of cells in the M1 immature stage (**Fig. 2H, Table S3**). After excluding significant differences in neutrophil maturation state across the traces, we next compared the transcriptomes for signatures that distinguish those derived from different HSPC traces. The results show that neutrophils derived from early- and mid-HSPCs upregulate metabolic processing of peroxide and NFkB-signaling, while those from late-HSPCs upregulated actin polymerization (**Fig. 2I, Table S3**). To further identify factors that may be driving distinct differentiation of neutrophils from the early-, mid- and late-HSPCs, we did pseudotime analysis using psupertime, a method that takes into account the real biological timepoints in pseudotime inference.^36^ The cytokine *il1b* appeared to be a driver of neutrophils differentiation from early-HSPCs but not from mid or late HPSCs, while *mmp9* was strongly associated with late-HSPCs (**Supplementary Fig. 3**). The finding further indicates that neutrophils arising from HSPCs at different developmental windows are transcriptionally distinct.

### HSPCs differentiate to ILC-like and Natural Killer-like cells

Initial analysis of the lymphoid subset of clusters in the scRNA-seq data revealed a highly heterogeneous pool (**Fig. 1B**). To date, rag+ T-cells are one of the only defined lymphocytes in young larval zebrafish, so the identity of the other lymphoid cells detected in the lineage traced scRNA-seq is unclear.^37,38^ Non-T lymphocytes like NK cells and ILCs were detected in juvenile and adult zebrafish but have not been detected in larval developmental stages.^28,39^ Mice and humans have a highly diverse repertoire of lymphoid cells emerging during embryogenesis and fetal life including many innate lymphocytes such as NK cells, ILC2s ILC3s, and B1A B-cells,^40–42^ thus we expected that some of the new populations could represent these cells. We posit that the enrichment of hematopoietic cells via *drl*-based lineage tracing and the isolation of mCherry+ cells found throughout the entire larvae could permit detection of rare immune cells, like ILCs, missed in prior larval zebrafish publications.^43,44^

Consistent with previous zebrafish work,^38^ we detected rag+ T lymphocytes expressing *lck, cd4-1, cd8b, tcra,* and *tcrb* in cluster 9 of the lymphoid lineage subset of cells in the scRNA-seq data (**Fig. 3A,B, Table S4**). Other less defined populations in larval zebrafish expressed top marker genes associated with mammalian ILCs and NK cells. Cluster 0 expresses a mix of ILC3-like markers such as *il7r, il23r, il17ra1a* and *tnfa*, as well as T-cell marker genes like *tcra* and *lck*. Cluster 2 expresses *eomesa*, *nkl.1, tbx21 (t-bet)*, and *gzm3.3* reminiscent of ILC1/NK-like cells. Cluster 3 expresses *il13, il4, gata3, mafa (MAF)* consistent with an ILC2-like identity. Cluster 7 highly expresses *gata3,* suggesting these could be ILC progenitors. Differential expression analysis between clusters 0, 2, 3, and 7, representative of the three groups of ILCs and potential progenitor, further highlights their unique transcriptional identities (**Fig. 3C, Table S4**).

**Figure 3.**
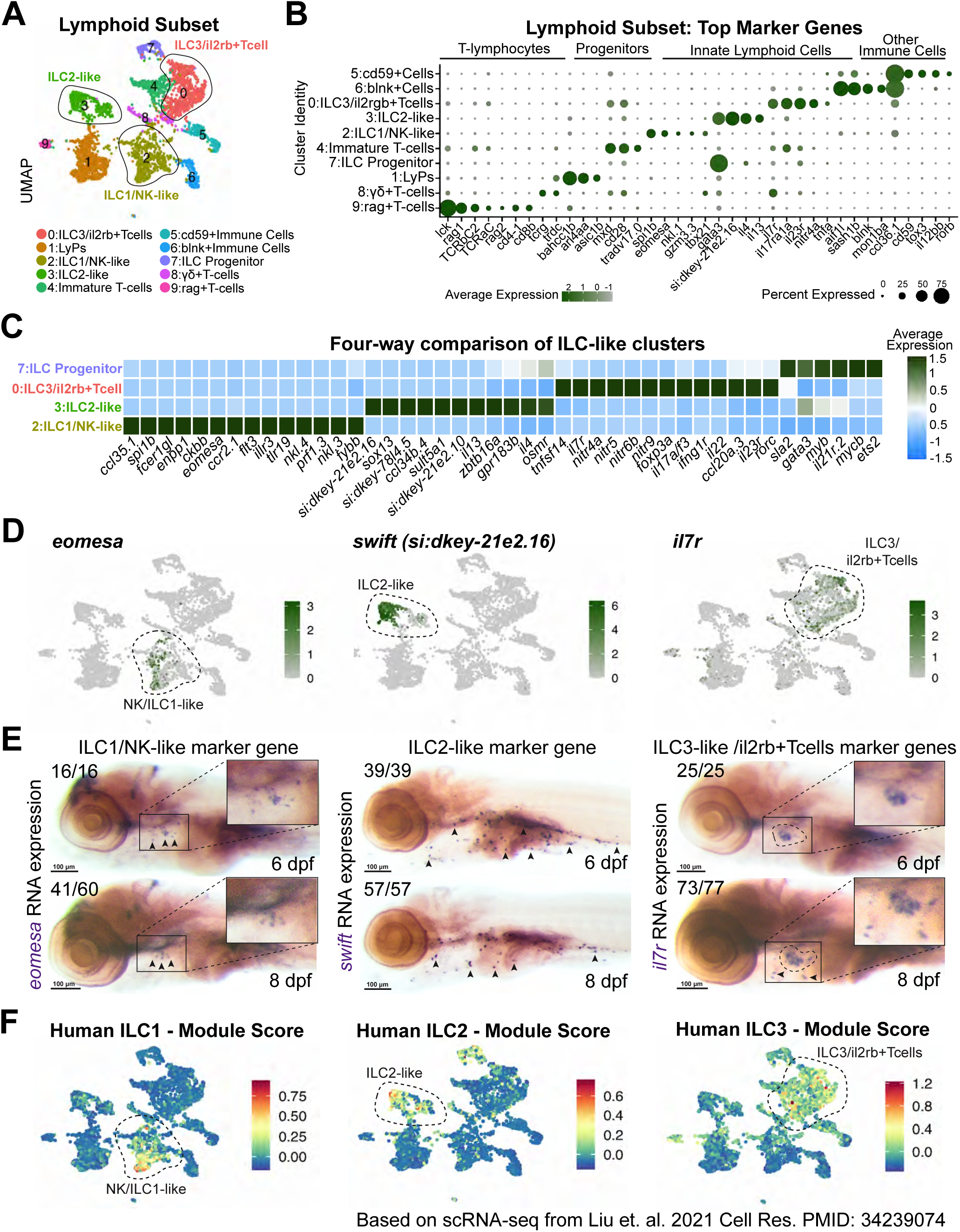
HSPCs differentiate to a variety of lymphoid lineage cells including ILC-like cells. (**A**) UMAP of lymphoid lineage subset with the three types of ILCs circled. (**B**) Dot plot showing select top marker genes of each cluster. (**C**) Heatmap of a four-way differential gene expression analysis comparing the putative ILC progenitor and ILC-like clusters. (**D**) Feature plot showing expression of three genes, *eomesa, si:dkey21e2.16, il7r*, selected for *in situ* hybridization analysis of the three ILC populations. (**E**) Images of RNA *in situ* hybridization for ILC marker genes at 6 (top) and 8 (bottom) dpf. Triangles point to quantified puncta and dotted lines circle the thymus. Numbers of images represent the number of larvae with similar phenotype over the total larvae tested. (**F**) Module scoring using human fetal ILC marker genes obtained from a scRNA-seq dataset.^46^

Aside from ILC-like identity, the lymphoid subset contains a variety of T-cells, progenitors, and other previously uncharacterized immune cells in larval zebrafish (**Fig. 3A,B**). Cluster 4 appears to be immature T cells, expressing *tcf7, mxd, cd28,* and *zap70*. Cluster 8 expresses *tcrg* and *trdc,* indicating a possible presence of gamma-delta T-cells. The putative identity of other clusters is less clear. For example, cluster 6 expresses *blnk*, *ccl36.1,* and *il12rb2* possibly indicating presence of a blnk+ B-cell progenitor although classic markers such as *pax5* are not detected (Table S4). Cluster 5 expresses genes like *il12bb, tox, cd59*, and *rorb*, that are expressed in many immune cell types making their identity less clear thus we refer to them as cd59+ Immune Cells.

The various ILC types have prototypic tissue locations. For example, mammalian ILC2s can be found residing in barrier tissues like lungs, intestine and skin.^45^ To determine if larval zebrafish ILC-like cells are in tissues that align with their hypothesized identity, we did whole mount *in situ* hybridization for top marker genes of ILC-like cells in 6 and 8 dpf larvae. We selected *eomesa* for the ILC1/NK-like cluster and *il7r* for the ILC3/il2rb+Tcells. To detect ILC2-like cells, we selected the gene *si:dkey21e2.16*, which is a predicted *<u>s</u>*erine-*t*ype endopeptidase, and named it *swift* (**Fig. 3D**). The expression of *eomesa* puncta is detectable around the thymic area, consistent with thymic NK cell localization^28^ (**Fig. 3E**). The expression of *swift* is detectable along the pharyngeal arches and intestines, which is consistent with mucosal localization of mammalian ILC2s. Lastly, *il7r* expression was localized to the thymus, but some puncta were present outside the thymus, along the pharyngeal arches. The two different locations are consistent with *il7r* marking multiple pots of cells, possibly ILC3s and il2rb+Tcells.

To determine if zebrafish ILCs are similar to human ILCs, we did gene module scoring using available datasets from human fetal stage ILCs.^46^ The gene sets enriched in human fetal ILCs were converted to zebrafish orthologs and then used to compute a score in the zebrafish lymphoid cell subset (**Fig. 3F, Table S4**). For each ILC subtype, the module scores were highest in the respective zebrafish ILC-like clusters. The result indicates that zebrafish ILCs are transcriptionally similar to the human counterparts. These data illustrate a previously unappreciated variety of lymphoid lineage immune cells that are detectable in hematopoietic and non-hematopoietic zebrafish tissues similar with their mammalian counterparts.

### Zebrafish ILC-like cells show similar dependencies on Il2rγ and Rag1 as mammalian ILCs

ILCs and adaptive T-lymphocytes, specifically *cd4+* T-helper cells, express similar genes.^19^ Therefore, identification of ILCs solely based on gene expression is inadequate to distinguish ILCs from T-helper cells. In humans and mice, both ILCs and T-lymphocytes depend on the function of interleukin 2 common gamma chain (IL2Rγ) because the immune cells express various interleukins that require IL2RG as a co-receptor.^47^ Unlike T-lymphocytes, ILCs lack antigen-specific receptors, and therefore do not require the function of Rag1 and Rag2 recombinase enzymes for their development.^20^ To better parse out ILCs from T-cells in larval zebrafish, we assessed the cells in *il2rga^Y91fs^*homozygous mutants^48^ and *rag1* mutagenized zebrafish. We anticipated that zebrafish ILC-like cells would be diminished in an *il2rga* mutant and unaffected by mutagenesis of *rag1*.

In zebrafish, Il2rγ is encoded by the duplicated genes *il2rga* and *il2rgb*, however only *il2rga* functions during developmental lymphopoiesis.^49^ We used the *il2rga^Y91fs^* zebrafish mutants to determine if larval ILCs require the function of Il2rγ. Heterozygous *il2rga^Y91fs^*fish containing either *Tg(drl:creER^T2^)* or *Tg(ubi:loxP-GFP-loxP-stop-mCherry)* were crossed to obtain *il2rga* wild-type, heterozygous, and homozygous embryos with both the tamoxifen-inducible Cre enzyme and the switch cassette (**Fig. 4A**). Early-HSPCs were labeled at 1 dpf and the mCherry expressing progeny were analyzed at 8 dpf using confocal imaging. In homozygous *il2rga^Y91fs^* mutants, mCherry-expressing cells were significantly reduced in the thymus and intestine compared with the wild-type and heterozygous siblings (**Fig. 4B**). However, mCherry-expressing cells in the CHT were unaffected by the loss of *il2rga,* consistent with the CHT mostly containing HSPC, myeloid, and erythroid cells^50^ which are not dependent on Il2rga. To further parse out if ILC-like cells are dependent on *il2rga*, we did *in situ* hybridization for *eomesa, swift,* and *il7r* at 6 dpf and showed reduced expression of all three markers in *il2rga^Y91fs^*homozygous mutants compared to wild-type and heterozygous siblings (**Fig. 4C**).

**Figure 4.**
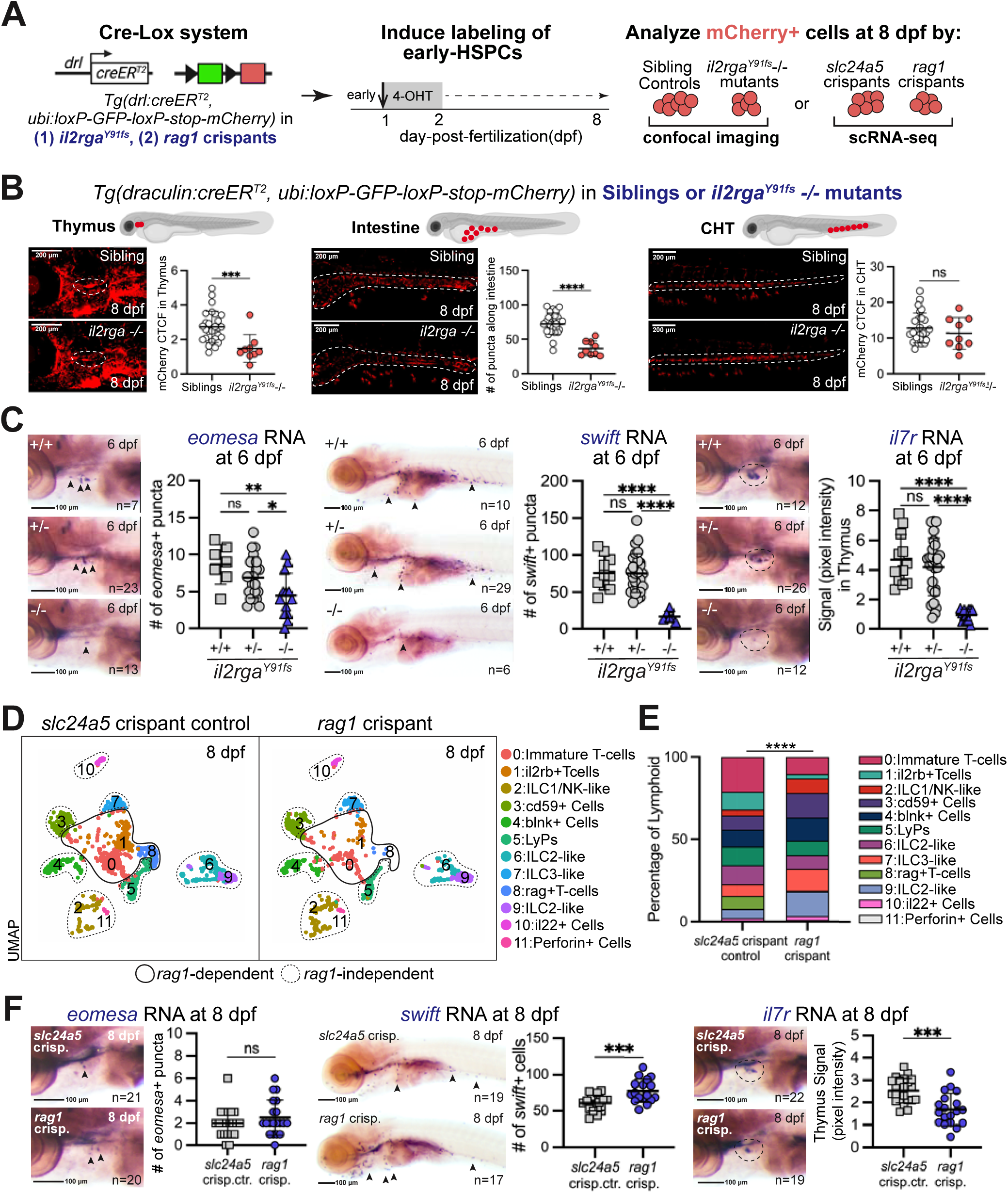
Zebrafish ILCs are Rag1-independent and Il2rγa-dependent. (**A**) Lineage tracing of early-HSPCs in *il2rga^Y91fs^* mutants or in *rag1* crispants at 8dpf. The *il2rga^Y91fs^* mutants were analyzed by confocal imaging. Crispants were subjected to scRNA-seq. The *slc24a5* (*golden*) locus was mutagenized as a control for the crispant experiment. (**B**) Confocal imaging of mCherry cells derived from early HSPCs in siblings and *il2rga^Y91fs^* homozygous mutants at 8 dpf (left). Graph showing quantification tissue-specific mCherry signal (right). Each point represents one fish. Significance computed with unpaired t-test. (**C**) Images of RNA *in situ* hybridization of *eomesa*, *swift*, and *il7r* (left). Graphs showing significant reduction of ILC markers in *il2rga^Y91fs^* homozygous mutants. Triangles point to quantified puncta and dotted lines circle the thymus. Each point represents one fish. Significance computed with one-way ANOVA with Tukey multiple testing corrections. (**D**) UMAP showing the lymphoid lineage subset of clusters in controls and *rag1* crispants with solid lines circling the *rag1*-dependent clusters and dotted lines circling *rag1-*independent clusters. (**E**) Stacked bar plot showing percentage of cells in each cluster relative to total lymphoid cells. Significance computed with chi-square test. See supplemental for absolute numbers. (**F**) RNA *in situ* hybridization for ILC marker genes at 8 dpf in controls and *rag1* crispants (left). Graphs of quantification of *eomesa, swift,* and *il7r* expression in the *rag1* crispants (right). Each point represents one fish. Significance computed with unpaired t-test. **p ≤0.01, ***p ≤0.001, **** p ≤0.0001

To distinguish ILCs from T-lymphocytes, we mutagenized the *rag1* locus using CRISPR/Cas9 in *Tg(drl:creER^T2^,ubi:Switch)* zebrafish larvae and then assessed T-lymphocytes and ILCs (**Supplementary Fig. 4A**). To control for off-target effects from mutagenesis, we compared *rag1* crispants to larvae mutagenized at the *slc24a5 golden* locus, a non-essential gene which is required for melanin pigmentation in zebrafish.^51^ In *rag1* crispants, T-cells were decreased when assessed by imaging of the T-cell fluorescent reporter *Tg(rag2:mCherry)* and by *in situ* hybridization for *rag1* and *tcra*, confirming decreased Rag1 functionality (**Supplementary Fig. 4B,C**).

To assess all lymphopoiesis in *rag1* and control *slc24a5* crispants, we performed scRNA-seq on the early-HSPC trace-derived mCherry-expressing cells isolated from 8 dpf larvae (**Fig. 4D, Supplementary Fig. 4D,E**). We hypothesized that Rag1-independent cells would show no change in abundance in the *rag1* crispants, while Rag1-dependent cells like T-lymphocytes would be diminished. As anticipated, cluster 8 identified as *rag*-expressing T-lymphocytes was diminished in *rag1* crispants (**Fig. 4D,E, Table S5**). Based on gene expression, clusters 1 and 7 appear to be the ILC3s and T-cells observed as cluster 0 in the original HSPC trace dataset (**Fig. 3A**), but due to the *rag1* genetic perturbation, the two cell types separated in this dataset (**Fig. 4D**). The T-cell identity of the newly discovered il2rb+ T cells (cluster 1) was confirmed by their decreased abundance in *rag1* crispants. Clusters 2, 6, 7, 9, and 11 were identified as NK or ILC-like based on top marker gene expression and were still present in high abundance in *rag1* crispants in accordance with ILCs not requiring the Rag enzyme for their development (**Fig. 4D,E, Supplementary Fig. 4D,E**). ILC2-like cells are represented by two clusters in this dataset, clusters 6 and 9. Cumulatively, the ILC2 levels were not significantly altered indicating Rag1 independence. Clusters 3 and 4 identified as cd59+ and blnk+ immune cells, respectively, were also Rag1-independent. *In situ* hybridization for markers of ILC1/NK-like and ILC2-like cells, *eomesa* and *swift* respectively, confirmed Rag1 independence (**Fig. 4F**). However, expression of *il7r* decreased in *rag1* crispants, indicating that the cells marked by *il7r* include T-lymphocytes consistent with the expression of *il7r* in both il2rb+ T-cell cluster 1 and ILC3-like cluster 7 (**Table S5**). In summary, transcriptionally detected ILC-like cells in zebrafish show genetic dependencies consistent with their proposed identity, and in agreement with mammalian ILCs.

### Zebrafish larval ILCs respond to viral mimicry

In adults, innate lymphocytes participate in the immune system response to pathogenic insults by viruses, bacteria, and helminths.^21^ Fetuses and newborns can also be exposed to these pathogens or their immunogenic products, yet the full immune system response during these life stages, especially by innate lymphocytes, is largely unknown. The identification of ILCs and the ability to look organism wide in larval zebrafish provides an opportunity to explore their early life immune responses. In humans, perinatal NK cells can respond to infectious agents such as single-stranded RNA viruses like influenza and SARS-CoV2.^52–54^ To test the responsiveness of the larval immune system to a relevant perinatal infection known to stimulate an innate lymphocyte population, we exposed larval zebrafish to the TLR7/8 agonist R848 (Resiquimod) as a means to mimic the presence of a ssRNA virus.^55^

To assess the global larval hematopoietic and immune response to viral ssRNA mimicry, we did whole body confocal imaging of early-HSPC-derived mCherry+ cells at 8 dpf following a two-day exposure to R848 (**Fig. 5A**). The treatment window of 6 – 8 dpf was selected as representative of a perinatal equivalent time point in zebrafish. At 8 dpf, the R848-treated larvae had a significant increase in mCherry signal in the thymus and CHT as well as an increased number of puncta in the intestine as compared to untreated controls (**Fig. 5B**).

**Figure 5.**
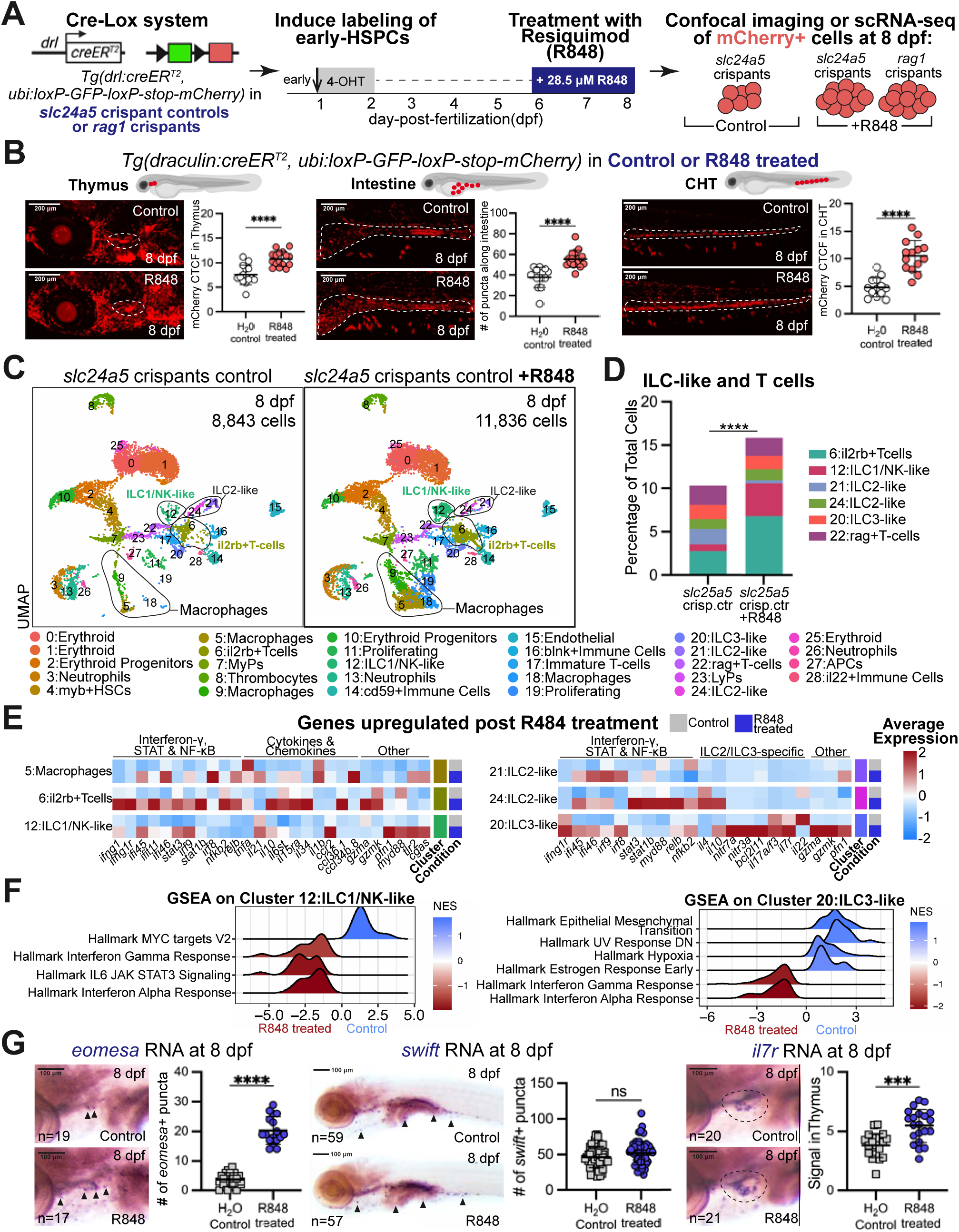
Viral mimicry elicits a response from the ILC-like cells in zebrafish larvae. **(A)** Early-HSPC trace-derived mCherry+ cells from controls and *rag1* crispants from either vehicle control or R848 treated groups were sequenced at 8 dpf. (**B**) Confocal imaging of mCherry+ cells derived from early HSPCs from 8 dpf controls and R848-treated larvae (left). Graphs depicting an increase in mCherry+ signal in the thymus, intestine and CHT of R848-treated larvae (right). Each point represents one fish. Significance computed with unpaired t-test. (**C**) UMAP showing all hematopoietic clusters detected at 8 dpf per condition with lines circling Macrophages, ILC1/NK-like, ILC2-like, and T-cells clusters that show the greatest response to the R848 agonist. (**D**) Bar plots showing the relative percentages of ILC/T-cell clusters in control and R848 treated groups out of all the cells shown in (C), Significance computed with chi-square test. See supplemental figure for absolute numbers. (**E**) Heatmaps showing average expression of select genes across the designated clusters in control and R848-treated groups. (**F**) GSEA analysis showing upregulation of interferon response genes in ILC clusters 12 and 20 in R848 treated larvae. (**G**) RNA *in situ* hybridization of *eomesa*, *swift,* and *il7r* with *eomesa* and *il7r* significantly increased in response to R848. Triangles point to quantified puncta and dotted lines circle the thymus. Each point represents one fish. Significance computed with unpaired t-test. ***p ≤0.001, **** p ≤0.0001

To determine which immune cells respond to the viral mimic, we did scRNA-seq on early-HSPC-derived mCherry-expressing cells from 8 dpf *slc24a5* crispant controls and *rag1* crispants treated with R848 (**Supplementary Fig. 5A,B**). The data were then used to decipher changes in cell number and gene expression as readouts of the response to viral ssRNA mimicry. Macrophages are known to respond strongly to TLR7/8 stimulation, so their response served as a control.^56^ In R848-treated larvae, there is a significant expansion of macrophage clusters 5, 9, and 18 (**Fig. 5C, Supplementary Fig. 5C**). Gene enrichment analysis of R848-induced transcriptional changes in these clusters revealed upregulation of genes that are concordant with metabolic changes and defense response activation (**Supplementary Fig. 5D, Table S5**).

Consistent with the work in human perinatal NK cells, there was an expansion of cluster 12 ILC1/NK-like cells in both *slc24a5* controls and in *rag1* crispants treated with R848 (**Fig. 5C,D, Supplementary Fig. 5C**). The cluster 6 il2rb+ T cells were also expanded in all R848-treated samples, but there was no change in the number of ILC2-like and ILC3-like cells. Upregulation in cytokine and immune signaling are another metric of an immune cell response, so we examined transcriptomic changes in T-cell and ILC populations in response to R848 exposure. In the cluster 6 il2rb+ T cells, there was an increase in genes associated with IFN-𝛾, STAT, and NF𝜅B signaling post R848 treatment (**Fig. 5E**). The cells also upregulated TNF𝛼, several cytokines, chemokines and granzymes, all consistent with an anti-viral immune response. The ILC1/NK-like cells showed an increase in *myd88,* IFNγ related genes like *ifng1 ifi46, ifi45, irf9*, STAT signaling components like *stat1b, stat3, il6st*, and higher expression of transcripts encoding cytokines like IL-10 and IL-21. Although we did not observe substantial changes in the number of ILC2-like and ILC3-like cells, these clusters also upregulated genes associated with IFN-𝛾, NFkB and STAT signaling implying they were responsive to the R848 stimulation (**Fig. 5E**). There was also some cell-type specific immune signaling upregulation, like ILC2-like cluster 24 had an increase in IL-4 (*il4*) and ILC3-like cluster 20 had an increase in IL-7R, IL-17A/F (*il7r, il17a/f3*), *nitr7a,* and *nitr3a* genes.

Gene set enrichment analysis (GSEA) using the hallmark gene set showed an Interferon response in ILC1/NK-like and ILC3-like cells treated with R848, and a STAT3 signaling enrichment in ILC1/NK-like cells (**Fig. 5F**). Enrichment analysis for GO terms (EnrichGO) revealed an upregulation of actin polymerization and motility GO terms in ILC1/NK-like cells which is consistent with a study showing increased migration of NK cells in responses to R848^57^(**Supplementary Fig. 5E, Table S5**). In agreement with this finding, we noted movement of mCherry+ lineage traced cells out of the thymus of R848-treated animals (**Supplementary Videos**).

As the scRNA-seq data suggested changes in cell number as well as potential mobility of ILC1/NK cells upon R848 exposure, we did *in situ* hybridization in whole larvae at 8 dpf for the ILC1/NK cell marker gene *eomesa*. (**Fig. 5G**). Concordant with the scRNA-seq result, *eomesa^+^* puncta were significantly increased by R848 treatment with more puncta noted outside the thymus along the pharyngeal arches in treated larvae compared to controls, suggesting that ILC1/NK-like cells are mobilized from the thymus. We also examined the expression of the ILC2 marker gene *swift* which was unchanged between controls and R848 treated (**Fig. 5G**). In agreement with the observed expansion of cluster 6 il2rb+Tcells in the sequencing data, expression of *il7r* was significantly increased in the thymus and appeared somewhat dispersed from this tissue in response to R848, suggestive of activation (**Fig. 5G**). Taken together, the evidence shows that R848 treatment elicited an immune response from the newly characterized ILC-like cells in larval zebrafish.

### Early-HSPCs preferentially contribute to adult tissue-resident lymphocytes

Temporal lineage tracing of larval hematopoiesis indicated that early HSPCs are the major contributors to larval lymphopoiesis, including ILCs (**Fig. 2**). Based on the strong preference of early HSPCs to generate lymphoid cells that seeded many tissues in larvae, we posited that these cells might be a significant source of adult tissue-resident lymphocytes. To address this question, we labeled early and late HSPCs using the *drl:CreERT;ubi:switch* system and then performed scRNA-seq on mCherry+ cells from the kidney marrow (equivalent to mammalian bone marrow), thymus, intestine, and tessellated lymphoid network (TLN, equivalent to skin immune cells)^58^ (**Fig. 6A**). The tissues were dissected from 4-month-old zebrafish and hematopoietic cells were isolated by fluorescent-activated cell sorting enriching for mCherry+ cells. The sequenced cells were analyzed in Seurat and clusters were identified based on top marker genes and the literature in the field. Across the tissues, we identified clusters representative of all expected hematopoietic cell types such as thrombocytes, HSCs, erythroid progenitors, neutrophils, macrophages, T-cells, B-cells, ILC1/NK-like, ILC2-like, ILC3-like, and dendritic cell populations (**Fig. 6B, Supplementary Fig. 6, Table 6**). In the lymphoid and barrier tissues, we also detected the new populations that we had found in larvae such as *blnk+* immune cells and *cd59+* immune cells. As expected, based on cell type distributions in mammalian counterparts and prior zebrafish studies,^28^ the kidney was enriched for myeloid lineage cells, while thymus, intestine and TLN contained more lymphoid lineage cells (**Fig. 6C**). The distribution of lymphoid cell types was not identical across the tissues. Consistent with mammalian bone marrow, the kidney marrow had mostly B-cells and T-cells with a lower abundance of ILC1/NKs, some ILC2-like and no ILC3-like cells^19^ (**Fig. 6B**). The intestine was enriched for T lymphocytes and ILC3-like cells but did not contain ILC1/NK-like populations. The TLN and the thymus both contained all three groups of ILC-like clusters.

**Figure 6.**
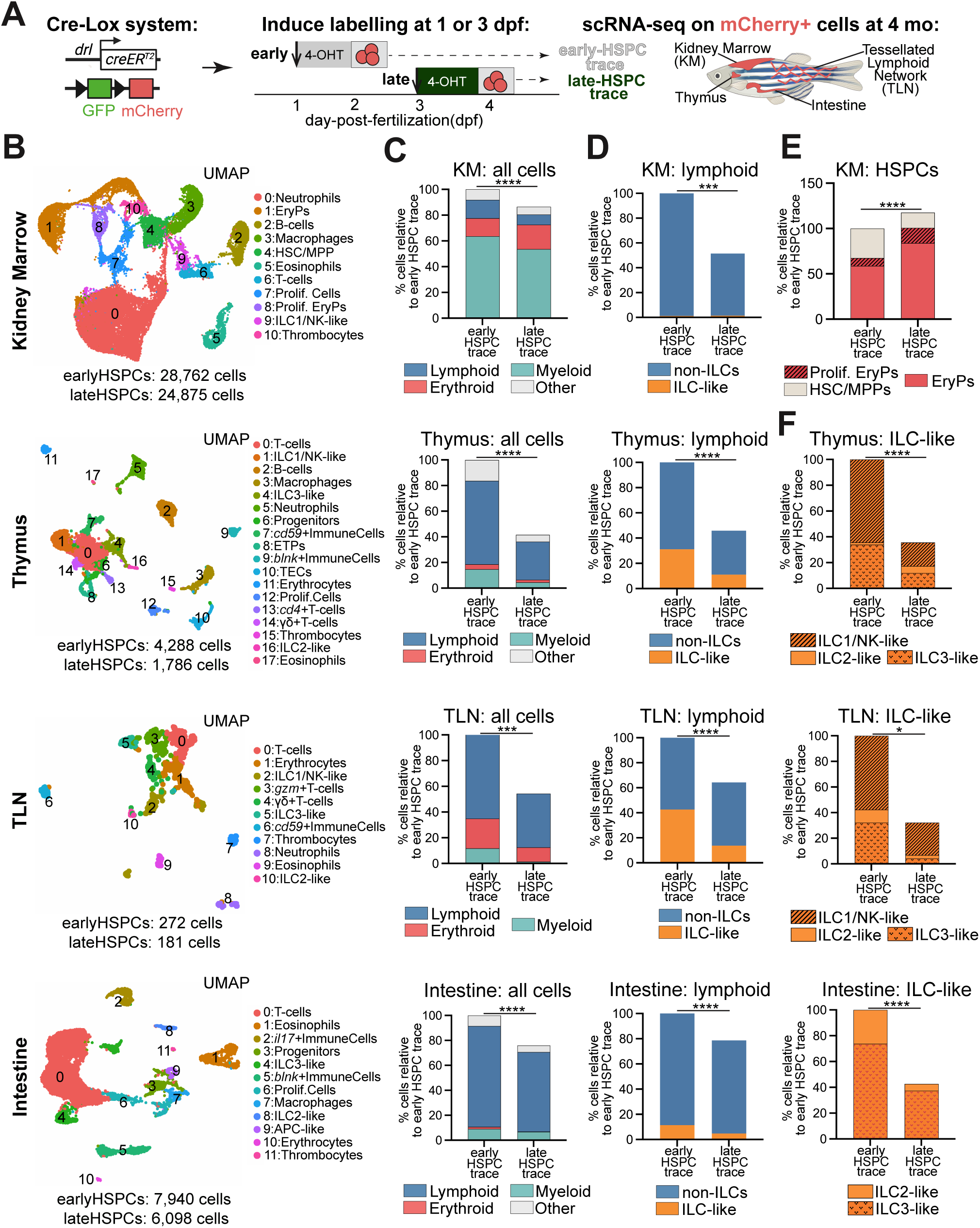
Early-arising HSPCs are the main contributors to adult tissue-resident lymphocytes. (**A**) Schema of early or late HSPC lineage tracing induced during embryonic development using the Cre-Lox inducible system followed by scRNA-seq of mCherry+ cells from kidney marrow, thymus, intestine and TLN isolated from 4 month adult zebrafish. (**B**) UMAPs showing cluster annotations per adult tissue with both traces combined. (**C**) Graphs showing percentage of hematopoietic cells across tissues relative to the early-HSPC trace. Significance computed with chi-square test. (**D**) Graphs showing percentage of lymphoid lineage cells across tissues relative to the early-HSPC trace, binned by ILC-like versus all other non-ILC lymphocytes. Significance computed with chi-square test. (**E**) Graph depicting percentage of HSCs and EryPs in kidney marrow relative to the early-HSPC trace. Significance computed with chi-square test. (**F**) Graphs depicting percentage of ILC-like cells across tissues relative to the early-HSPC trace. Significance computed with chi-square test. *p<0.05, ***p ≤0.001, ****p ≤0.0001

To facilitate cross trace comparisons of early *versus* late HSPC contributions to adult hematopoietic and immune landscapes, we normalized the data across samples by organ. Each organ has a unique abundance of immune cells, and our goal is to define which fraction of those are derived from early *versus* late HSPCs. For similar comparisons in mammalian studies, the absolute number of hematopoietic and immune cells within a tissue of interest are quantified first using pan-hematopoietic antibodies and then the relative fraction arising from a given lineage trace is calculated. Due to the lack of antibodies or transgenic lines in zebrafish that label all hematopoietic and immune cells, we adopted a slightly different approach. As the number of immune cells detected in all tissues analyzed was highest in the early HSPC trace samples, we normalized the abundance of immune cells for both samples to the early HSPC trace for each organ. The analysis revealed early HSPCs contribute about two-fold more to the hematopoietic and immune compartments in the thymus and skin-TLN (**Fig. 6C,D**). There is also a greater fraction of early HSPC-derived cells in the kidney marrow and intestine, although to a lesser extent.

Examination of the relative distribution of hematopoietic and immune cells in these adult tissues derived from the early *versus* late HSPCs revealed similar differentiation preferences as observed in larval life. For example, similar to the observation during larval stages, late-HSPC-traced cells in the kidney marrow seemed biased towards the erythroid lineage as evidence by more erythroid progenitors derived from the late-traced sample (**Fig. 6C&E**). In contrast, early HSPCs contributed more to the lymphoid compartment in the kidney marrow, thymus, TLN and to a lesser extent the intestine (**Fig. 6D**).

The developmental origin of barrier lymphoid immunity, especially ILCs, is poorly understood, so we next examined the contribution of early *versus* late HSPCs to specific lymphoid cell populations within the skin-TLN and intestine. In the skin-TLN, there was a 3.1-fold difference between early and late traces especially the ILC1/NK-like and ILC3-like cells (**Fig. 6F**). The trend was similar in the intestine with 2.5-fold more ILC-like cells derived from early HSPCs than late, particularly the ILC3 subtype (**Fig. 6F**). These findings indicate that although there are distinct distributions of lymphoid cells across barrier tissues, ILC-like cells found in them are mostly derived from HSPCs arising earlier in development. Overall, these results suggests that the developmental bias to lymphopoiesis in early-arising HSPCs and erythropoiesis in late-arising HSPCs persists not only to adult hematopoietic tissues like the kidney and thymus but also key barrier immune organs like the skin and intestine.

### Developmental time imparts HSPCs with distinct transcriptional signatures and Runx1 sensitivities

Given the observation that early-arising and late-arising HSPCs show persistent differentiation preferences (**Fig. 6**), we posited that there could be different transcriptional signatures in HSPCs from the early and late traces that hint at their final lineage biases. Cluster 4 within the kidney marrow samples expresses genes representative of HSCs and multipotent progenitors (MPPs), thus we examined the differentially expressed genes between the early-and late-derived cluster 4 HSC/MPPs to determine if the time of embryonic origination imparted a long-lasting transcriptional signature to adult HSPCs. Early-HSPC trace derived HSC/MPPs expressed higher levels of lymphoid and immune-related genes like *socs1a* and *hes6* and while HSC/MPPs from the late-HSPC trace expressed higher levels of erythroid lineage genes like *cahz* and *alas2* (**Fig. 7A**). KM HSC/MPPs derived from early-traced HSPCs expressed higher levels of xenobiotic response genes while those from late HSPCs show enriched expression of genes involved in erythrocyte and myeloid lineage differentiation (**Fig. 7B, Table S7**).

**Figure 7.**
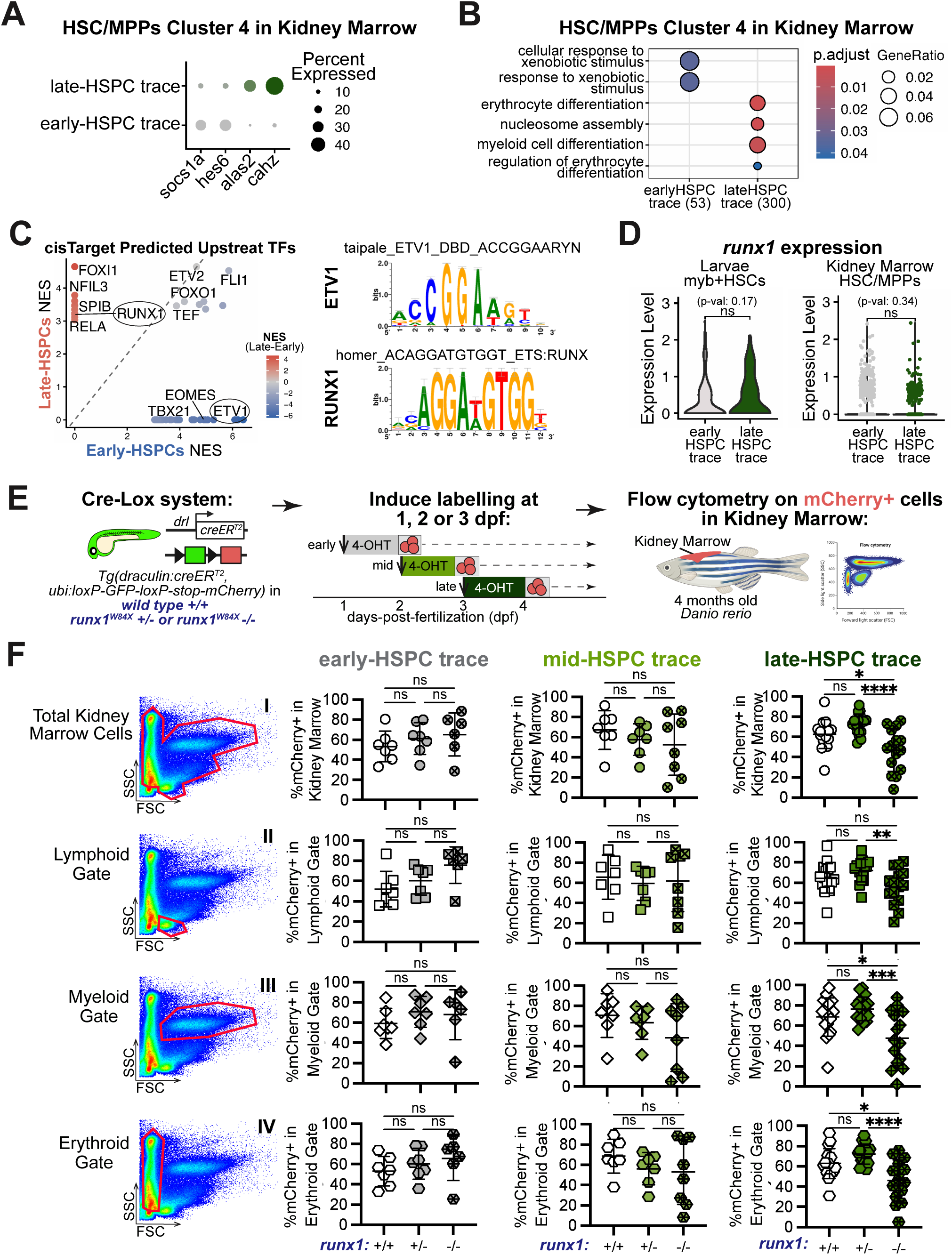
Early and late HSPCs show unique sensitivities to the transcription factor Runx1. (**A**) Dot plot showing the expression of select lymphoid and erythroid lineage genes in early and late HSPC trace-derived kidney marrow cluster 4 HSC/MPPs. (**B**) EnrichGO Biological Processes pathway enrichment analysis comparing kidney marrow HSC/MPPs between the two traces. (**C**) Scatter plot showing the predicted upstream transcription factors regulating early and late HSPCs based on upstream motif analysis of genes differentially expressed between early and late trace-derived kidney marrow cluster 4 HSC/MPP. The RUNX1 (late) and ETV1 (early) motifs are highlighted as examples of putative trace-specific upstream regulators. (**D**) Violin plots showing the expression of *runx1* in larval and adult HSC clusters in early and late HSPC traces. Significance computed with a Wilcoxon test. (**E**) Lineage tracing in wild-type, heterozygous and homozygous *runx1^W84X^* mutants following embryonic early, mid, and late HSPC labeling. Kidney marrow cells from 4-month-old zebrafish were analyzed for percentage of early, mid, or late trace-derived mCherry+ cells by flow cytometry. (**F**) Flow plots showing the lineage(s) examined: I- total kidney marrow; II- Lymphoid gate; III- Myeloid gate; IV- Erythroid gate. The full flow strategy is shown in Figure S7. Graphs quantifying the percentage of mCherry+ cells from the three traces across *runx1* genotypes. Each point represents one fish. Significance computed with one-way ANOVA with Tukey multiple testing corrections. *p<0.05, **p ≤0.01, ***p ≤0.001, **** p ≤0.0001

Next, we searched for upstream transcription factors (TFs) that might impart the transcriptional differences that distinguish early and late arising HSPCs. We analyzed TF motifs enriched across the genes differentially expressed between KM HSC/MPPs derived from early-*versus* late-HSPC traces, which revealed several predicted upstream TFs with trace-specific enrichment (**Fig. 7C, Table S7**). Among the high confidence TFs, ETV1, an ETS family transcription factor linked to neurogenesis, was early-trace specific, while RUNX1, a Runt-family transcription heavily involved in hematopoiesis, was late-trace specific. These data hint at differential TF activity in the early and late HSPC traces. We also inspected expression of these TFs in the KM HSC/MPP cluster as well as in the larval *myb+* HSC cluster to assess if they displayed trace-specific gene expression. The zebrafish *etv1* gene is not expressed in our larvae datasets, and *runx1* expression levels were similar in both traces at larval and adult stages (**Fig. 7D**).

These findings led to the hypothesis that early *versus* late arising HSPCs could be differentially sensitive to Runx1 levels. RUNX1 is a transcription factor critical for HSPC formation.^59,60^ In mice, *Runx1* loss is embryonic lethal in large part due to the lack of hematopoietic progenitors during fetal development.^59,61^ In zebrafish, some *runx1* homozygous loss-of-function mutants die due to hematopoietic failure but some can survive to adulthood because of genetic compensatory mechanisms that seem to restore most of the hematopoietic system.^62,63^ To decipher if HSPCs forming during different developmental windows are differentially sensitive to Runx1 levels, we did lineage tracing of early, mid, and late HSPCs in *runx1^w84x^* wild-type, heterozygous, and homozygous mutants and then used flow cytometric quantification to examine the relative frequency of mCherry+ cells within all marrow cells and within each blood lineage of 4 month old zebrafish (**Fig. 7E, Supplementary Fig. 7**). The early and mid HSPCs contributed a similar percentage of mCherry+ cells to the total kidney marrow across the genotypes (**Fig. 7F**). However, late HSPCs showed a reduction in the percentage of mCherry cells in the *runx1^W84X^* homozygous mutant. The data suggests that loss of Runx1 impacts the late HSPCs ability to contribute to adult hematopoiesis in the kidney marrow. Furthermore, compared to wild-type controls, the late HSPCs in *runx1* homozygous mutants contributed significantly less to the myeloid and erythroid lineages, while the lymphoid lineage was not significantly affected (**Fig. 7F**). Combined, the data suggests that Runx1 loss may be more severely impairing erythroid-myeloid biased HSCs that are labeled by the late trace in our system compared to the lymphoid-biased cells labeled by the early trace.

## DISCUSSION

In this study, we uncovered two major aspects of developmental hematopoiesis. We illustrated that long-lasting lineage bias is etched into HSPCs at distinct developmental windows with lymphoid lineage bias preceding other differentiation preferences, insights important for understanding origins of clonal hematopoiesis and age-related hematologic health risks. Furthermore, our characterization of innate lymphocytes in the larvae presents a new frontier for zebrafish immunology with implications on understanding lymphocyte ontogeny and perinatal lymphoid immune responses.

While hematopoietic transplantation is the gold standard for examining HSPC functionality, seminal studies measuring native hematopoiesis revealed greater complexity and functionality within the HSPC pool.^15,64^ For example, studies examining native hematopoiesis in murine embryos illustrated that progenitors arising before definitive HSCs can differentiate to adaptive lymphocytes like T and B cells that persist to adulthood.^5,17^ Furthermore using transposon-based lineage tracing, Patel *et al.* revealed that a subset of later arising HSCs are biased toward generating megakaryocytes.^5^ In zebrafish, Tian *et al.* and He *et al.* showed that the progenitor generating the earliest wave of thymic seeding T-cells was HSC independent.^23,24^ Our lineage-tracing findings build on these works by establishing that embryonic lymphoid-producing early HSPCs can generate a wide variety of adaptive and innate lymphocytes during larval life (**Fig. 1-4**). Furthermore, we demonstrated that HSPCs arising earlier during embryogenesis contribute to the life-long pool of lymphocytes residing in barrier tissues (**Fig. 6**). Prior work showed tissue resident macrophages arose from an earlier HSC-independent progenitor wave,^16,65^ and our data now supports a broader contribution of early embryonic HSPCs to tissue resident lymphoid cells. Moreover, in a similar vein to Patel *et al.*, we observed that later arising HSPCs in zebrafish show a distinct lineage preference to those emerging earlier in embryonic development. The subset of HSPCs captured by our later lineage tracing paradigm showed a bias toward erythropoiesis in larvae and adults (**Fig. 1, 2, 6**).

Mechanistically, regulation of divergent HSPCs and their differentiation output remains to be elucidated. Recent work showed that HSPC heterogeneity is initiated in the hemogenic endothelium and is influenced by factors like apico-basal polarity of cells^66^, and by Notch and Wnt signaling whose regulation in the embryo affects adulthood lineages.^67^ For our work, we posited that transcriptional wiring imparted during development would leave hallmarks in adult HSPCs. Upstream regulator analysis in adult cells uncovered RUNX1 signatures in the late-derived HSPCs. Our assessment of hematopoiesis in zebrafish *runx1* mutants supports the transcriptional observation and illustrates that late-derived HSPCs are more dependent on Runx1 (**Fig. 7**). The effect was not due to level of *runx1* expression but rather differential functional requirement. Studies in mice also find temporal requirements for hematopoietic transcription factors C-MYB^68^ and RUNX1^69^. Dependency on C-MYB is used to distinguish early primitive and later definitive waves of hematopoiesis,^16,65,68^ and RUNX1 dependency can distinguish early from late yolk sac EMPs.^69^ Given the embryonic lethality of homozygous murine mutants, experiments connecting embryonic HSPCs and adult hematopoiesis remains limited. As zebrafish can survive for weeks without blood due to their external development,^32^ they offer an opportunity to reveal novel insights on hematopoietic ontogeny, including our finding that distinct adult-contributing HSPCs display differential Runx1 sensitivity. Future mechanistic studies are needed to determine how Runx1 is imparting lineage bias on HSPCs during embryogenesis.

The finding of an array of ILCs located across many hematopoietic and non-hematopoietic depots throughout young larvae begs the question of why an organism would produce ILCs so early in development. Perhaps the earlier wave is needed as a line of defense against pathogens before the adaptive immune system matures in the fetus. Consistent with a few limited studies on fetal NK cells following pathogen exposure,^53,54^ our results show that in addition to NK cells, ILC-like populations can respond to viral-like stimulation early in larval life.

To understand if or how ILC ontogeny can affect their function, the embryonic origins of the multitude of tissue resident ILCs need better resolution.^24,70^ In larval zebrafish, we find that ILC3s are produced by the earlier-arising HSPCs (**Fig. 2**), which is similar to the developmental origins of murine lymphoid tissue inducers, which are a subtype of ILC3s.^71,72^ The other ILC subtypes are not as well resolved in the larvae, but examination of the adult tissue resident ILCs shows that a majority of ILCs and NK cells identified in the thymus, skin-TLN, and intestine are derived from early-arising HSPCs (**Fig. 6**). The mechanism underlying how and when these ILCs enter these tissues is unclear but work from Wang *et al.* showing that murine embryonic ILC progenitors seed the liver, lung, and intestine during fetal development and later differentiate into distinct ILC subsets suggest the pattern is established early in life.^73^

Overall, our work illustrates that HSPC lineage preference manifests in a temporal manner during embryogenesis with lymphoid bias arising first. Moreover, we illuminated that despite divergence in tissue distribution, tissue resident lymphocytes across many adult tissues are derived from the lymphoid-biased early arising HSPC population. Understanding the regulation of this temporal embryonic imprint has the potential to uncover a connection between HSPC ontogeny to a variety of human autoimmune disorders and leukemias as well as age-associated disorders linked to HSPC heterogeneity.

## Acknowledgements

T.V.B. was supported by grants from the National Institutes of Health (NIH) R01DK121738, R01DK131445, R01DK141169, and the Edward P. Evans Foundation. AN was supported by NIH F31HL167600, NYSTEM award C34874GG (PI Frenette), and a Liang Zhu Memorial Fellowship. BAU was supported by NIH F31HL152562 and T32GM007288 (PI Akabas), KJ was supported by NIH F31HL180025, T32GM145438 (PI Query), and R01HL153920 (PI Zheng). Instrumental feedback to the development of this project was provided by Dr. Kira Gritsman, Dr. Ulrich Steidl, and Dr. Meelad Dawlady. We thank the Langenau lab for sharing *il2rga^Y91fs^* mutant lines generated as part of R24OD016761. We thank the core facilities instrumental for this work that are supported by the Einstein National Cancer Institute’s grant number P30CA013330, including: Flow Cytometry Core Facility at Einstein and core members Yu (Joey) Zhang and Ming Liu for assistance with FACS; the Einstein Genomics Core and core member David Reynolds for scRNA library preparation; and the Analytical Imaging Facility. We thank Dr. Roshan Sharma for meaningful feedback on scRNA-seq data and Dr. Eliseo Castillo and Dr. Keir Balla for fruitful discussions on innate lymphoid cells. Thank you to Dr. Noura Ghazale for meaningful discussions and technical assistance, and Dr. Kathryn Potts for meaningful review and discussion of the manuscript. Graphics were created in BioRender with copyright https://BioRender.com/rsdc34d and https://BioRender.com/riwx66c

## Author Contributions

Conceptualization, TVB, AN, BAU; Methodology, TVB, AN, BAU, KJ, DZ; Investigation, TVB, AN, BAU; Writing—Original Draft, TVB, AN; Writing—Review & Editing, TVB, AN, BAU, KJ, DZ; Funding Acquisition, TVB, AN; Supervision, DZ, TVB.

## METHODS

### EXPERIMENTAL MODEL AND STUDY PARTICIPANT DETAILS

Zebrafish were bred and maintained as described previously.^74^ All fish were maintained according to Institutional Animal Care and Use Committee (IACUC)-approved protocols in accordance with the Albert Einstein College of Medicine research guidelines. Sex determination in zebrafish is complex and cannot be accounted for at larval stages,^75^ but for all adult stages we used equal numbers of males and females.

### Transgenic and Mutant Lines

For lineage tracing, we used *Tg(drl:creER^T2^)* and the *Tg(ubi:loxP-GFP-loxP-stop-mCherry)* also known as *Tg(ubi:Switch).*^25,26^ For all experiments, fish with two copies of *Tg(drl:creER^T2^,ubi:Siwtch)* transgenes were crossed to WT AB fish (from ZIRC) so all resulting experimental animals were hemizygous. We used the published *runx1^W84X^*mutants and *il2rga^Y91fs^* mutants.^48,60^ Genotyping of *runx1^W84X^* mutants was done using a qPCR with a custom designed Taqman SNP probe GGTTCTGGTACCTTGAAGGCGATGGGCAGGGTCTTGTTGCAGCG[C/T]CAGTGTGTCGGCAGGACGGAGCAGAGGAAGTTCGGGCTGTCGG from Life Technologies. The *il2rga^Y91fs^* genotyping was done as described in Tang *et al*.^48^ For lineage tracing in *runx1^W84X^* mutants, we crossed *Tg(drl:creER^T2^),runx1^W84X^+/- to Tg(ubi:Switch),runx1^W84X^+/-* zebrafish. To validate the functional consequences of *rag1* crispants on T-cells, we used the *Tg(rag2:mCherry).*^76^

### Lineage Tracing Assay

To induce fluorescent-based lineage tracing of HSPCs, embryos containing *Tg(drl:creER^T2^,ubi:Siwtch)* were incubated in a 28°C incubator with 12 𝜇M 4-hydroxytamoxifen (4-OHT) (Sigma H7904) for 20 hours, starting at 30 hpf for early HSPCs, at 54 hpf for mid HSPCs, and at 86 hpf for late HSPCs. Hydroxytamoxifen was washed out three times with embryo water then larvae were kept in a clean dish until larval endpoint or raised in our zebrafish aquatic system until adult time points.

### Single Cell RNA Sequencing

#### Cell isolation and Library Preparation

##### Larvae

For the early-HSPCs and mid-HSPCs cohorts, 200-250 larvae were used at one-day-post labelling, 6 dpf, and 10 dpf. To obtain enough fluorescent cells from the late-HSPCs derived cohort at the three timepoints (4, 6, and 10 dpf), we processed 1,100, 200, and 170 zebrafish larvae, respectively. For the *rag1* crispants/R848 treatment experiment, 35 8 dpf larvae were processed per condition. For all samples, whole larvae were minced with a razor blade and digested with a 1:65 dilution of a 5 mg/mL liberase (Sigma 5401127001) stock for 15 minutes at 37°C, followed by FBS inactivation as previously described.^22^ The mCherry-expressing cells were sorted into 500 𝜇L IMDM + 10% FBS buffer using the BD FACSAria II instrument at room temperature. The cells were immediately processed for cell capture by the Einstein genomics facility, using the Chromium Next GEM Single Cell 3ʹ Kit v3.1 for samples in Figures 1-3 and the v4 kit for *rag1* crispant/R848 samples in Figures 4 & 5. Cell numbers for each experiment are listed in Tables S1-5.

##### Adult Tissues

For kidney, intestine, thymus, and Tessellated Lymphoid Network (TLN) experiments, two adult fish were used per condition, one male and one female. All four organs were collected from the same fish. Fish were euthanized with tricaine, and TLNs were dissected first, then thymus, intestines, and kidneys last. TLN was dissected according to described protocols.^58^ In brief, a razor blade was used to scrape off the scales into 1X DPBS, and the resulting suspension was filtered to remove the scales. Cells were centrifuged and resuspended in DPBS for FACS. For preparing intestinal tissue, we developed a protocol based on two publications.^77,78^ Animals were not fed for 24 hours prior to dissection to improve the cleanliness of the intestinal tissue. Intestines were dissected from the animals and then minced with scissors in cold DPBS. The tissue was then digested with 1 mg/mL Collagenase IV (ThermoFisher 17104019) for 30 minutes at 37°C. The digested cells were filtered through a 40 µm cell strainer, centrifuged, and resuspended in DPBS with 2 mM EDTA. Kidneys were dissected and then mechanically triturated in FACS buffer to release the cells into suspension.^79^ FACS buffer contains 0.9XD-PBS, 5% FBS, and 1% Pen/Strep. To enrich for myeloid and lymphoid lineages in the kidney, we dissected kidneys and then lysed the erythrocytes using a 1:10 dilution of ACK buffer for 10 minutes. Cells were washed with 1XDPBS, filtered, and resuspended in FACS buffer. Thymus was dissected under a fluorescent microscope and dissociated by trituration in 500 µL HBSS (No Calcium, No Magnesium) with 0.5%BSA + DAPI and filtered through a 40 µm cell strainer. All FACS sorting was done on the BD FACS Aria II machine and mCherry positive cells were sorted into 500 𝜇L IMDM + 10% FBS. Libraries were prepared immediately by the genomics core at Einstein using the Chromium Next GEM Single Cell 3ʹ Kit v4. The libraries were sequenced on the Illumina NovaSeq 6000 machine. For early-HSPCs condition, we sequenced 33K, 35K, 86K, 33K reads per cells for kidney, TLN, intestine, and thymus respectively. For late-HSPCs condition we sequenced 42K, 49K, 130K, 51K reads per cell for kidney, TLN, intestine, and thymus respectively. Cell numbers for each experiment are listed in Tables S6.

#### Quality Control and Analysis

##### Larvae datasets

Fastq files were processed in Cell Ranger 7.0.1, except the *rag1* crispant/R848 experiment which was done in version 8.0.1. To generate the counts matrix in Cell Ranger, we made a custom reference genome based on *Danio rerio* GRCz11 with the addition of mCherry, GFP, and T cell specific genes: TCRaC, TCRbC1 and TCRbC2. All subsequent dimension reduction and clustering was done in Seurat version 5.2.0.^29^ No batch effect was apparent between samples, so analysis was done without batch corrections (Figure S1-2). Cells with less than 200 genes and more than 10% mitochondrial DNA were filtered out (Figure S1,5,6). For the lineage tracing dataset in Figure 1, we used the 4,000 top variable features for initial dimensional reduction, with 30 principal components (PC) and at cluster resolution of 1. After non-hematopoietic cells were removed, the remaining hematopoietic cells were re-analyzed with top 2,000 variable features, 30 PCs and at cluster resolution of 1 (Figure S2A). Two more lower quality clusters were identified and removed to obtain the final clustering using 2,000 variable features, 20 PCs, and cluster resolution of 1. For the lymphoid only subset analysis in Figure 3, we used 2,000 variable genes, 25 PCs, and 0.3 cluster resolution. For *rag1* crispant/R848 dataset in Figures 4-5, initial clustering was done using 4,000 variable genes, 30 PCs, and cluster resolution of 1. After removing non-hematopoietic cells, final clustering was done using 2,000 variable genes, 20 PCs, and a cluster resolution of 1. For analysis of the lymphoid only subset in Figure 4, we used 1,000 variable genes, 15 PCs, and 0.2 cluster resolution.

##### Adult datasets

All FASTQ files were processed in Cell Ranger 8.0.1 using the default counts setting and the custom-built *Danio rerio* GRCz11 genome. All clustering analysis was done in Seurat version 5.2.0.^29^ We filtered low quality cells using the 10% mitochondrial genes threshold and filtered out cells containing less than 200 genes. To filter out possible doublets, cells containing a high number of genes were filtered out using a different cutoff for each sample. For kidney samples, initial clustering revealed a batch effect among the neutrophil clusters, so we did a batch correction method on the kidney sample. We used anchor based CCA integration followed by clustering using 20 Principal Components (PCs), and 0.5 cluster resolution. Clusters 18 and 19 were removed from analysis because they were an unclear mix of cells. For thymus samples, no batch effects were present, so clustering was done using 1,500 variable features, 20 PCs and cluster resolution of 0.5. For TLN, initial clustering was done using 4,000 variable features, 20 PCs and a 0.5 cluster resolution. The dataset contained many erythrocytes, so we further subset away the erythroid populations to examine only the lymphoid and myeloid immune cells. The final clustering for TLN was done using 2,700 variable features, 20 PCs and 0.7 cluster resolution. For intestine, initial clustering revealed batch effects and many ribosomal protein genes across clusters. We removed all genes starting with *rps, rpl, rsl* and did anchor based CCA integration to correct for batch effect. Final clustering was done using 20 PCs and 0.5 resolution, and 4 small clusters were removed due to unclear mix of cells.

#### Cluster identification

For all cluster identification, we used a combination of literature-based marker genes and an assessment of top marker genes using the FindAllMarkers function in Seurat which compares the target cluster to other clusters using the Wilcoxon Rank Sum test. Module scoring using the built in AddModuleScore() function was also used to aid in cluster identification.

#### Differential Expression and GO Enrichment Analysis

For differential gene expression analysis, two clusters of interest were compared using the FindMarkers function in Seurat using standard settings for Wilcoxon Rank Sum test. For 4-way gene expression comparison, clusters were subset and compared using the FindMarkers function with standard settings. For gene enrichment analysis, the list of genes obtained from differential expression analysis was filtered to include only those with adjusted p-value of less that 0.05, and an average fold-change greater than 1.5. Using the enrichGO function available in ClusterProfiler package,^80^ we identified overrepresented GO terms that belong to the Biological Processes category. Displayed in Figure 2I, 7B, Supplemental Figure 2D, 5D,E are selected GO categories from the output list with the full lists found in Supplemental Tables S2, S3, S5, S7. The GSEA analysis for Hallmark gene set in Figures 1I and 5F were done by fold-change ranking.

#### Pseudotime Analysis

The software Psupertime^36^ was used to temporally order each cluster of cells in the hematopoietic atlas dataset from the different lineage traces. We include in the supplement results from psupertime on neutrophils (Figure S3). Psupertime was run with default settings with the exception of “min_expression” set to 0.00 instead of 0.01 to include genes that were expressed in any cell rather than at least 1% to ensure that rare lineage genes were not missed due to low abundance of some cell types in some traces.

#### Upstream Transcription Factor and Motif Enrichment

Upstream motif analysis was done using the RcisTarget. The list of zebrafish genes that are differentially expressed between early- and late-HSPC trace derived adult kidney marrow HSC/MPP was filtered by log fold-change greater than 0.6 and adjusted p-value less than 0.05. Genes were converted to human orthologs using gProfiler. For motif rankings, we used the hg38 file with 10kb up and down of TSS. Displayed in Figure 7C are the predicted high confidence transcription factors with NES greater than 3.0. The full list of predicted upstream regulators is found in Table S7.

### Confocal Microscopy and Analysis

The CSU-W1 Nikon spinning disc confocal was used for all high-resolution imaging of lineage tracing in wild-type and mutant conditions and in drug-treated larvae. For live imaging, larvae were anesthetized with 0.01% tricaine and mounted in 4% methylcellulose on glass bottom dished. Images were acquired using 10x lens with the following parameters: 600 millisecond exposure, and 561𝜆 laser at 65% power and the z-stack was set to 6 𝜇m size steps. The videos were generated by imaging 6dpf larvae embedded in 1% low melt agarose and anesthetized with tricaine. Larvae were mounted after 1 hour of R848 treatment, and then R848 was re-added after mounting during image acquisition. Images were acquired for 4 hours total, with 4-minute loops and a step size of 2.5 𝜇m. Images were analyzed in Fiji (ImageJ) version 2.1.0.^81^ To calculate mCherry signal in the thymus and CHT, we used Corrected Total Cell Fluorescence formular: (Integrated density of region of interest) - (Area of region of interest x Mean fluorescence of 6 different background regions). All quantification of live imaging was done blindly to genotype and/or condition, with the exception of images in Figure 1G and 2A.

### *In situ* Hybridization

The *tcra* RNA probe was made from Addgene plasmid #11262 gifted by Lisa Steiner.^82^ The *il7r* RNA probe was made from a PCR template that was amplified from cDNA collected from 10 dpf whole larvae. The following primers were used to generate the PCR template: (1) TAATACGACTCACTATAGGGGGCCTTCCTTTCACTTTCAG and (2) CATTAACCCTCACTAAAGGGAACATGGATTAATGTCTATAGCAGAGGA. Transcription of the anti-sense *il7r* RNA was done using T3 RNA polymerase (Sigma 11031163001). To make *eomesa* and *swift* probes, the DNA sequences were synthesized and put into pTWIST-kan high copy vector by TWIST Bioscience. These probes are available from Addgene [*eomesa-#*236391*; swift- #*236392; *il7r-* 236393]. For the *eomesa* probe, a 971-bp fragment containing exon 7 through the 3’UTR region and flanked by T7 and T3 primers was synthesized. For the *swift* probe, a 922-bp fragment was made containing the entire coding region of the gene. Gene-specific primers that contained T3 RNA polymerase sequence (forward) and T7 RNA polymerase sequence (reverse) at the 5’end were then used to generate a linear PCR-amplified template for *in vitro* transcription using T7 RNA polymerase (Sigma 10881775001) and DIG RNA Labeling Mix (Sigma 11277073910). The whole mount RNA *in situ* hybridization was done based on established zebrafish protocols with modifications for older larvae.^83^ For 6 & 8 dpf stages, pigment was removed by bleaching with a solution of 0.8%KOH, 0.9% H_2_O_2_, and 0.1% Tween20 for 35 & 45 min, respectively. Proteinase K (Sigma 3115887001) working concentration was increased to 15 mg/mL, and digestion time was extended to 35 & 40 min for 6 & 8 dpf, respectively. After digestion, samples were fixed with 0.25% glutaraldehyde for 25 and 30 minutes for 6 & 8 dpf, respectively. Incubation in the RNA probes was done for two nights at 70°C to maximize detection. To visualize the RNA probe signal, we used anti-DIG (Sigma 11093274910) followed by colorimetric detection using NBT/BCIP (Fisher PR-S3771). Samples were incubated in NBT/BCIP for 1-3 days for *eomesa* or *il7r*, and 8 hours for *swift,* all at room temperature. For experiments with *il2rga^Y91fs^* mutants, each embryo was imaged, quantified, and then genotyped.

### CRISPR/Cas9 Mutagenesis

The crispant approach was used to mutagenize the *rag1* locus, where three guide RNAs and S.p. Cas9 Nuclease V3 protein (IDT 1081058) were assembled into a complex and injected into one-cell stage embryos.^84^ Each guide RNA mix was assembled by annealing a tracr RNA (IDT 1072532) with the locus specific crRNA for 5 min at 95°C, and cooled at room temperature for 10 min. Three guides were used to target the *rag1* locus, two of them were designed in CHOPCHOP^85^ (*rag1* guide #1 AATGATGCCCACATCCCAGG, guide#2: AATGATGCCCACATCCCAGG), and one sequence from ZFIN (*rag1* guide#3: GGACTGCTGCCATAAGGGGA, ZDB-CRISPR-201209-4) (Figure S4). As a control for mutagenesis, an established *slc26a4* (golden) guide RNA was used: GGACAGACGTGTTTCTCCAG.^86^ Each single guide RNA was assembled into a complex with Cas9 separately. For the control, *slc24a5* targeting complex alone was done, and for *rag1* crispants, 3 *rag1* guides + 1 *slc24a5* guide were mixed in equal volumes to make a four-guide complex. A T7 endonuclease assay (NEB M0302L) was used to confirm mutagenesis, where cutting occurs only in the presence of heteroduplex DNA indicating mutagenesis (Figure S4).

### Resiquimod R848 Drug Treatment

The 500 𝜇g of R848 powder was obtained from InvivoGen (tlrl-r848-1) and reconstituted in 500 𝜇L of water for a final concentration of 1 𝜇g/𝜇L or 2.85 mM. To treat larvae, the stock was diluted in embryo water to a working concentration of 28.5 𝜇M. Larvae were incubated with R848 starting at 6 dpf and analyzed at 8 dpf.

### Flow Cytometry

Adult 4 month old zebrafish were euthanized on ice and kidneys were dissected and placed into 500 𝜇L of FACS buffer containing 1µg/mL DAPI (Sigma D8417). The FACS buffer is composed of 0.9XD-PBS, 5% FBS, and 1% Pen/Strep. To release cells into suspension, the dissected tissue was gently triturated with a p1000 pipette about 20 times. The cell suspension was filtered through a 40 µm cell strainer and collected into flow tubes by centrifugation at 2500 rpm for 40 seconds. For results shown in Figure 7, the red blood cells were not lysed, and samples were assessed on LSRII (BD Bioscience) instrument. Voltages were set appropriately and kept consistent between replicates. Analysis of data was done in FlowJo v10.8.1 software and gates were set as shown in Supplemental Figure 7. A wild-type control sample was used to delineate the fluorescent from non-fluorescent cells.

### QUANTIFICATION AND STATISTICAL ANALYSIS

Analysis was done in GraphPad Prism 10.2.3. For all experiments with three or more groups, a one-way ANOVA with Tukey multiple testing corrections was used. For two groups, an unpaired t-test was used as indicated in the figure legends. A Chi-square test was used to compare number of cells per cluster in the stacked bar plots. In all plots, one point is a single zebrafish larva or adult. For single-cell RNA sequencing, one replicate was done per condition. Confocal imaging experiments were done with at least three independent experiments, and *in situ* hybridization experiments were done at least two independent times. All error bars show standard deviation from the mean. In all figures: ns = not significant, * - p< 0.05, ** - p ≤0.01, *** - p≤0.001, **** - p≤0.0001.

## Material Availability

Reagents used in this study will be made available upon request. Plasmids for making RNA probes are deposited at Addgene IDs 236391, 236392, 236393.

## Data and Code Availability

The sequencing data is available via the GEO accession number: GSE292726 The processed data can be viewed at: https://sites.google.com/view/tvbowmanlab/developmental-hematopoiesis-atlas Codes can be made available upon request.

## REFERENCES

1 Muller-Sieburg, C. E., Cho, R. H., Karlsson, L., Huang, J. F. & Sieburg, H. B. Myeloid-biased hematopoietic stem cells have extensive self-renewal capacity but generate diminished lymphoid progeny with impaired IL-7 responsiveness. Blood 103, 4111–4118 (2004). 10.1182/blood-2003-10-3448

2 Challen, G. A., Boles, N. C., Chambers, S. M. & Goodell, M. A. Distinct hematopoietic stem cell subtypes are differentially regulated by TGF-beta1. Cell Stem Cell 6, 265–278 (2010). 10.1016/j.stem.2010.02.002

3 Sieburg, H. B. et al. The hematopoietic stem compartment consists of a limited number of discrete stem cell subsets. Blood 107, 2311–2316 (2006). 10.1182/blood-2005-07-2970

4 Ross, J. B. et al. Depleting myeloid-biased haematopoietic stem cells rejuvenates aged immunity. Nature 628, 162–170 (2024). 10.1038/s41586-024-07238-x

5 Patel, S. H. et al. Lifelong multilineage contribution by embryonic-born blood progenitors. Nature 606, 747–753 (2022). 10.1038/s41586-022-04804-z

6 Gore, A. V., Pillay, L. M., Venero Galanternik, M. & Weinstein, B. M. The zebrafish: A fintastic model for hematopoietic development and disease. Wiley Interdiscip Rev Dev Biol 7, e312 (2018). 10.1002/wdev.312

7 Kobayashi, M. & Yoshimoto, M. Multiple waves of fetal-derived immune cells constitute adult immune system. Immunol Rev 315, 11–30 (2023). 10.1111/imr.13192

8 Jagannathan-Bogdan, M. & Zon, L. I. Hematopoiesis. Development 140, 2463–2467 (2013). 10.1242/dev.083147

9 Canu, G. & Ruhrberg, C. First blood: the endothelial origins of hematopoietic progenitors. Angiogenesis 24, 199–211 (2021). 10.1007/s10456-021-09783-9

10 McGrath, K. E., Frame, J. M. & Palis, J. Early hematopoiesis and macrophage development. Semin Immunol 27, 379–387 (2015). 10.1016/j.smim.2016.03.013

11 Le Guyader, D. et al. Origins and unconventional behavior of neutrophils in developing zebrafish. Blood 111, 132–141 (2008). 10.1182/blood-2007-06-095398

12 Dege, C. et al. Potently Cytotoxic Natural Killer Cells Initially Emerge from Erythro-Myeloid Progenitors during Mammalian Development. Dev Cell 53, 229–239 e227 (2020). 10.1016/j.devcel.2020.02.016

13 Yoshimoto, M. et al. Embryonic day 9 yolk sac and intra-embryonic hemogenic endothelium independently generate a B-1 and marginal zone progenitor lacking B-2 potential. Proc Natl Acad Sci U S A 108, 1468–1473 (2011). 10.1073/pnas.1015841108

14 Iturri, L. et al. Megakaryocyte production is sustained by direct differentiation from erythromyeloid progenitors in the yolk sac until midgestation. Immunity 54, 1433–1446 e1435 (2021). 10.1016/j.immuni.2021.04.026

15 Ginhoux, F. et al. Fate mapping analysis reveals that adult microglia derive from primitive macrophages. Science 330, 841–845 (2010). 10.1126/science.1194637

16 Schulz, C. et al. A lineage of myeloid cells independent of Myb and hematopoietic stem cells. Science 336, 86–90 (2012). 10.1126/science.1219179

17 Kobayashi, M. et al. HSC-independent definitive hematopoiesis persists into adult life. Cell Rep 42, 112239 (2023). 10.1016/j.celrep.2023.112239

18 Neo, W. H., Lie, A. L. M., Fadlullah, M. Z. H. & Lacaud, G. Contributions of Embryonic HSC-Independent Hematopoiesis to Organogenesis and the Adult Hematopoietic System. Front Cell Dev Biol 9, 631699 (2021). 10.3389/fcell.2021.631699

19 Cherrier, D. E., Serafini, N. & Di Santo, J. P. Innate Lymphoid Cell Development: A T Cell Perspective. Immunity 48, 1091–1103 (2018). 10.1016/j.immuni.2018.05.010

20 Spits, H. et al. Innate lymphoid cells--a proposal for uniform nomenclature. Nat Rev Immunol 13, 145–149 (2013). 10.1038/nri3365

21 Yuan, T. et al. Innate lymphoid cells and infectious diseases. Innate Immun 30, 120–135 (2024). 10.1177/17534259241287311

22 Ulloa, B. A. et al. Definitive hematopoietic stem cells minimally contribute to embryonic hematopoiesis. Cell Rep 36, 109703 (2021). 10.1016/j.celrep.2021.109703

23 Tian, Y. et al. The first wave of T lymphopoiesis in zebrafish arises from aorta endothelium independent of hematopoietic stem cells. J Exp Med 214, 3347–3360 (2017). 10.1084/jem.20170488

24 He, S. et al. In vivo single-cell lineage tracing in zebrafish using high-resolution infrared laser-mediated gene induction microscopy. Elife 9 (2020). 10.7554/eLife.52024

25 Mosimann, C. et al. Ubiquitous transgene expression and Cre-based recombination driven by the ubiquitin promoter in zebrafish. Development 138, 169–177 (2011). 10.1242/dev.059345

26 Mosimann, C. et al. Chamber identity programs drive early functional partitioning of the heart. Nat Commun 6, 8146 (2015). 10.1038/ncomms9146

27 Henninger, J. et al. Clonal fate mapping quantifies the number of haematopoietic stem cells that arise during development. Nat Cell Biol 19, 17–27 (2017). 10.1038/ncb3444

28 Rubin, S. A. et al. Single-cell analyses reveal early thymic progenitors and pre-B cells in zebrafish. J Exp Med 219 (2022). 10.1084/jem.20220038

29 Hao, Y. et al. Dictionary learning for integrative, multimodal and scalable single-cell analysis. Nat Biotechnol 42, 293–304 (2024). 10.1038/s41587-023-01767-y

30 Ouyang, J. F., Kamaraj, U. S., Cao, E. Y. & Rackham, O. J. L. ShinyCell: simple and sharable visualization of single-cell gene expression data. Bioinformatics 37, 3374–3376 (2021). 10.1093/bioinformatics/btab209

31 Murayama, E. et al. Tracing hematopoietic precursor migration to successive hematopoietic organs during zebrafish development. Immunity 25, 963–975 (2006). 10.1016/j.immuni.2006.10.015

32 Soza-Ried, C., Hess, I., Netuschil, N., Schorpp, M. & Boehm, T. Essential role of c-myb in definitive hematopoiesis is evolutionarily conserved. Proc Natl Acad Sci U S A 107, 17304–17308 (2010). 10.1073/pnas.1004640107

33 Gao, X., Xu, C., Asada, N. & Frenette, P. S. The hematopoietic stem cell niche: from embryo to adult. Development 145 (2018). 10.1242/dev.139691

34 Ginhoux, F. & Guilliams, M. Tissue-Resident Macrophage Ontogeny and Homeostasis. Immunity 44, 439–449 (2016). 10.1016/j.immuni.2016.02.024

35 Kirchberger, S. et al. Comparative transcriptomics coupled to developmental grading via transgenic zebrafish reporter strains identifies conserved features in neutrophil maturation. Nature Communications 15, 1792 (2024). 10.1038/s41467-024-45802-1

36 Macnair, W., Gupta, R. & Claassen, M. psupertime: supervised pseudotime analysis for time-series single-cell RNA-seq data. Bioinformatics 38, i290–i298 (2022). 10.1093/bioinformatics/btac227

37 Langenau, D. M. et al. In vivo tracking of T cell development, ablation, and engraftment in transgenic zebrafish. Proc Natl Acad Sci U S A 101, 7369–7374 (2004). 10.1073/pnas.0402248101

38 Langenau, D. M. & Zon, L. I. The zebrafish: a new model of T-cell and thymic development. Nat Rev Immunol 5, 307–317 (2005). 10.1038/nri1590

39 Hernandez, P. P. et al. Single-cell transcriptional analysis reveals ILC-like cells in zebrafish. Sci Immunol 3 (2018). 10.1126/sciimmunol.aau5265

40 Dorshkind, K. & Crooks, G. Layered immune system development in mice and humans. Immunol Rev 315, 5–10 (2023). 10.1111/imr.13198

41 Tang, Y. et al. Emergence of NK-cell progenitors and functionally competent NK-cell lineage subsets in the early mouse embryo. Blood 120, 63–75 (2012). 10.1182/blood-2011-02-337980

42 Schneider, C. et al. Tissue-Resident Group 2 Innate Lymphoid Cells Differentiate by Layered Ontogeny and In Situ Perinatal Priming. Immunity 50, 1425–1438 e1425 (2019). 10.1016/j.immuni.2019.04.019

43 Petrie-Hanson, L., Hohn, C. & Hanson, L. Characterization of rag1 mutant zebrafish leukocytes. BMC Immunol 10, 8 (2009). 10.1186/1471-2172-10-8

44 Yoder, J. A. et al. Developmental and tissue-specific expression of NITRs. Immunogenetics 62, 117–122 (2010). 10.1007/s00251-009-0416-5

45 Naito, M. & Kumanogoh, A. Group 2 innate lymphoid cells and their surrounding environment. Inflamm Regen 43, 21 (2023). 10.1186/s41232-023-00272-8

46 Liu, C. et al. Delineating spatiotemporal and hierarchical development of human fetal innate lymphoid cells. Cell Res 31, 1106–1122 (2021). 10.1038/s41422-021-00529-2

47 Colonna, M. Innate Lymphoid Cells: Diversity, Plasticity, and Unique Functions in Immunity. Immunity 48, 1104–1117 (2018). 10.1016/j.immuni.2018.05.013

48 Tang, Q. et al. Dissecting hematopoietic and renal cell heterogeneity in adult zebrafish at single-cell resolution using RNA sequencing. J Exp Med 214, 2875–2887 (2017). 10.1084/jem.20170976

49 Sertori, R. et al. Conserved IL-2Rgammac Signaling Mediates Lymphopoiesis in Zebrafish. J Immunol 196, 135–143 (2016). 10.4049/jimmunol.1403060

50 Xia, J. et al. A single-cell resolution developmental atlas of hematopoietic stem and progenitor cell expansion in zebrafish. Proc Natl Acad Sci U S A 118 (2021). 10.1073/pnas.2015748118

51 Lamason, R. L. et al. SLC24A5, a putative cation exchanger, affects pigmentation in zebrafish and humans. Science 310, 1782–1786 (2005). 10.1126/science.1116238

52 Veneziani, I., Alicata, C., Moretta, L. & Maggi, E. Human toll-like receptor 8 (TLR8) in NK cells: Implication for cancer immunotherapy. Immunol Lett 261, 13–16 (2023). 10.1016/j.imlet.2023.07.003

53 Gee, S. et al. The legacy of maternal SARS-CoV-2 infection on the immunology of the neonate. Nat Immunol 22, 1490–1502 (2021). 10.1038/s41590-021-01049-2

54 Li, J. et al. Impaired NK cell antiviral cytokine response against influenza virus in small-for-gestational-age neonates. Cell Mol Immunol 10, 437–443 (2013). 10.1038/cmi.2013.31

55 Veneziani, I. et al. Toll-like receptor 8 agonists improve NK-cell function primarily targeting CD56(bright)CD16(-) subset. J Immunother Cancer 10 (2022). 10.1136/jitc-2021-003385

56 Lopez-Cuevas, P. et al. Reprogramming macrophages with R848-loaded artificial protocells to modulate skin and skeletal wound healing. J Cell Sci 137 (2024). 10.1242/jcs.262202

57 Zhou, J., Nouri-Shirazi, M., Tang, H., Shen, Y. & Zeng, M. R848/TLR7-Mediated Stronger CD8+ T Immunity Is Dependent on DC-NK Cell Interactions. Int Arch Allergy Immunol 183, 860–875 (2022). 10.1159/000522364

58 Robertson, T. F. et al. A tessellated lymphoid network provides whole-body T cell surveillance in zebrafish. Proc Natl Acad Sci U S A 120, e2301137120 (2023). 10.1073/pnas.2301137120

59 Chen, M. J., Yokomizo, T., Zeigler, B. M., Dzierzak, E. & Speck, N. A. Runx1 is required for the endothelial to haematopoietic cell transition but not thereafter. Nature 457, 887–891 (2009). 10.1038/nature07619

60 Jin, H. et al. Definitive hematopoietic stem/progenitor cells manifest distinct differentiation output in the zebrafish VDA and PBI. Development 136, 647–654 (2009). 10.1242/dev.029637

61 Chen, M. J. et al. Erythroid/myeloid progenitors and hematopoietic stem cells originate from distinct populations of endothelial cells. Cell Stem Cell 9, 541–552 (2011). 10.1016/j.stem.2011.10.003

62 Sood, R. et al. Development of multilineage adult hematopoiesis in the zebrafish with a runx1 truncation mutation. Blood 115, 2806–2809 (2010). 10.1182/blood-2009-08-236729

63 Bresciani, E. et al. Redundant mechanisms driven independently by RUNX1 and GATA2 for hematopoietic development. Blood Adv 5, 4949–4962 (2021). 10.1182/bloodadvances.2020003969

64 Sun, J. et al. Clonal dynamics of native haematopoiesis. Nature 514, 322–327 (2014). 10.1038/nature13824

65 Gomez Perdiguero, E. et al. Tissue-resident macrophages originate from yolk-sac-derived erythro-myeloid progenitors. Nature 518, 547–551 (2015). 10.1038/nature13989

66 Torcq, L., Vivier, C., Schmutz, S., Loe-Mie, Y. & Schmidt, A. A. Single-cell and in situ spatial analyses reveal the diversity of newly born hematopoietic stem cells and of their niches. Development 152 (2025). 10.1242/dev.204454

67 Ghersi, J. J. et al. Haematopoietic stem and progenitor cell heterogeneity is inherited from the embryonic endothelium. Nat Cell Biol 25, 1135–1145 (2023). 10.1038/s41556-023-01187-9

68 Tober, J., McGrath, K. E. & Palis, J. Primitive erythropoiesis and megakaryopoiesis in the yolk sac are independent of c-myb. Blood 111, 2636–2639 (2008). 10.1182/blood-2007-11-124685

69 Tober, J., Yzaguirre, A. D., Piwarzyk, E. & Speck, N. A. Distinct temporal requirements for Runx1 in hematopoietic progenitors and stem cells. Development 140, 3765–3776 (2013). 10.1242/dev.094961

70 Dignum, T. et al. Multipotent progenitors and hematopoietic stem cells arise independently from hemogenic endothelium in the mouse embryo. Cell Rep 36, 109675 (2021). 10.1016/j.celrep.2021.109675

71 Simic, M. et al. Distinct Waves from the Hemogenic Endothelium Give Rise to Layered Lymphoid Tissue Inducer Cell Ontogeny. Cell Rep 32, 108004 (2020). 10.1016/j.celrep.2020.108004

72 Elsaid, R. et al. A wave of bipotent T/ILC-restricted progenitors shapes the embryonic thymus microenvironment in a time-dependent manner. Blood 137, 1024–1036 (2021). 10.1182/blood.2020006779

73 Wang, X. et al. Innate lymphoid cells originate from fetal liver-derived tissue-resident progenitors. Sci Immunol 10, eadu7962 (2025). 10.1126/sciimmunol.adu7962

74 Lawrence, C. in Methods in Cell Biology Vol. 104 (eds H. William Detrich, Monte Westerfield, & Leonard I. Zon) 429–451 (Academic Press, 2011).

75 Aharon, D. & Marlow, F. L. Sexual determination in zebrafish. Cell Mol Life Sci 79, 8 (2021). 10.1007/s00018-021-04066-4

76 Harrold, I. et al. Efficient transgenesis mediated by pigmentation rescue in zebrafish. Biotechniques 60, 13–20 (2016). 10.2144/000114368

77 Jones, L. O. et al. Single-cell resolution of the adult zebrafish intestine under conventional conditions and in response to an acute Vibrio cholerae infection. Cell Rep 42, 113407 (2023). 10.1016/j.celrep.2023.113407

78 Jiang, M. et al. Characterization of the Zebrafish Cell Landscape at Single-Cell Resolution. Front Cell Dev Biol 9, 743421 (2021). 10.3389/fcell.2021.743421

79 Mahony, C. B. & Monteiro, R. Protocol for the analysis of hematopoietic lineages in the whole kidney marrow of adult zebrafish. STAR Protoc 5, 102810 (2024). 10.1016/j.xpro.2023.102810

80 Yu, G., Wang, L. G., Han, Y. & He, Q. Y. clusterProfiler: an R package for comparing biological themes among gene clusters. OMICS 16, 284–287 (2012). 10.1089/omi.2011.0118

81 Schindelin, J. et al. Fiji: an open-source platform for biological-image analysis. Nat Methods 9, 676–682 (2012). 10.1038/nmeth.2019

82 Danilova, N. et al. T cells and the thymus in developing zebrafish. Dev Comp Immunol 28, 755–767 (2004). 10.1016/j.dci.2003.12.003

83 Thisse, C. & Thisse, B. High-resolution in situ hybridization to whole-mount zebrafish embryos. Nat Protoc 3, 59–69 (2008). 10.1038/nprot.2007.514

84 Burger, A. et al. Maximizing mutagenesis with solubilized CRISPR-Cas9 ribonucleoprotein complexes. Development 143, 2025–2037 (2016). 10.1242/dev.134809

85 Labun, K. et al. CHOPCHOP v3: expanding the CRISPR web toolbox beyond genome editing. Nucleic Acids Res 47, W171–W174 (2019). 10.1093/nar/gkz365

86 Avagyan, S. et al. Resistance to inflammation underlies enhanced fitness in clonal hematopoiesis. Science 374, 768–772 (2021). 10.1126/science.aba9304

