## Supplementary Figures and Legends for "Developmental Timing Establishes Hematopoietic Stem and Progenitor Cell Lineage Bias"

### SUPPLEMENTAL FIGURES AND FIGURE LEGENDS

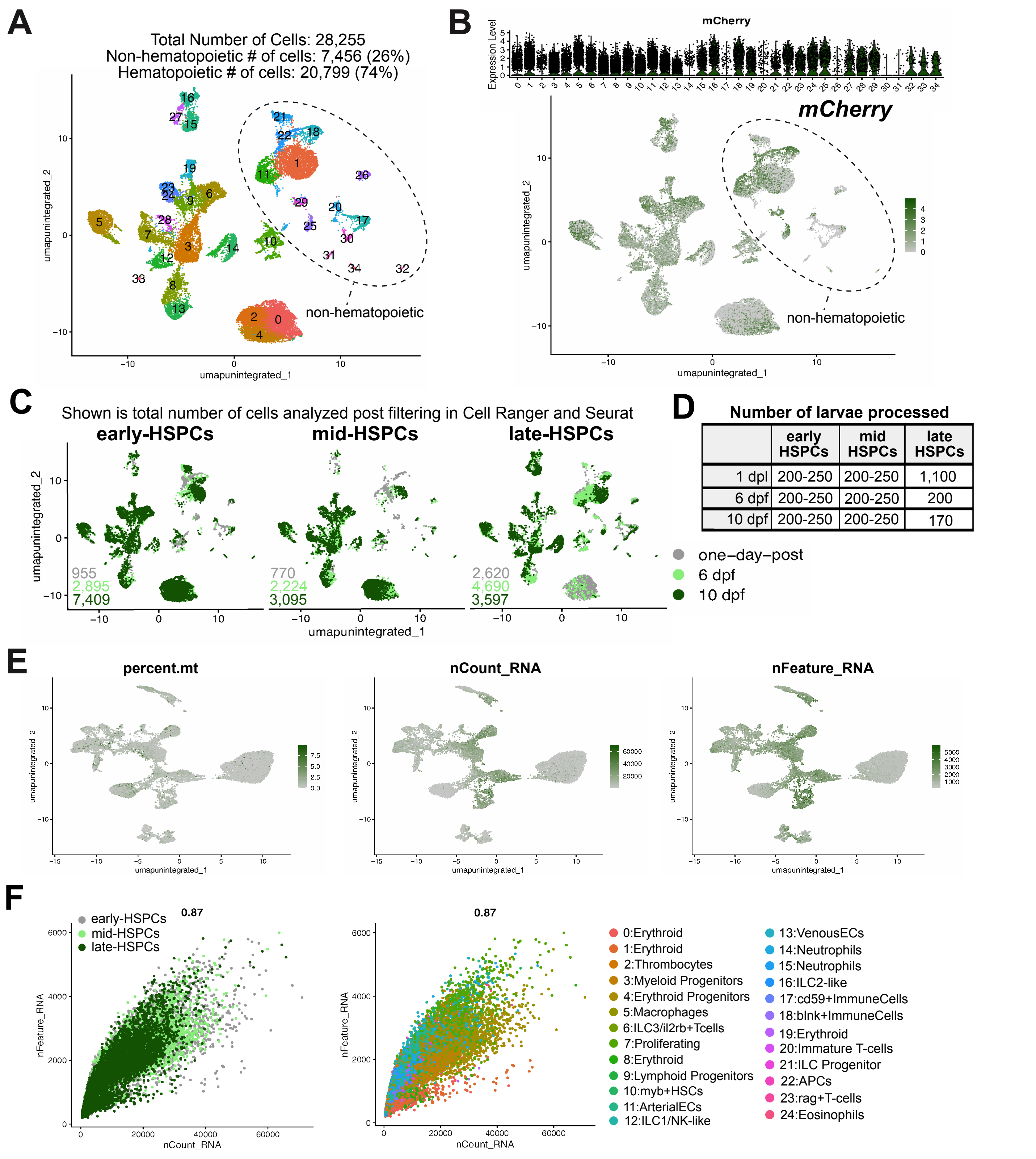


| | early HSPCs | mid HSPCs | late HSPCs |
|---|---|---|---|
| 1 dpl | 200-250 | 200-250 | 1,100 |
| 6 dpf | 200-250 | 200-250 | 200 |
| 10 dpf | 200-250 | 200-250 | 170 |

**Supplemental Figure 1. Filtering and quality control.** (A) UMAP of 34 clusters representing all the cells detected in early-, mid-, and late-HSPC trace scRNA-seq. Non-hematopoietic populations excluded from downstream analysis are circled. (B) Violin plots (top) and feature plot (bottom) of mCherry transcript in all populations. (C) UMAP of all cells split by trace and developmental timepoint illustrating no batch effects between conditions. (D) Table with number of larvae used for each timepoint and trace. (E) Feature plots of hematopoietic and endothelial cell clusters showing percent mitochondria genes (left) ranging from 0-10%, number of counts (middle), and number of features or genes (right) expressed in each cell and cluster. (F) Scatter plots of counts versus features by HSPC trace (left) and by cluster (right).

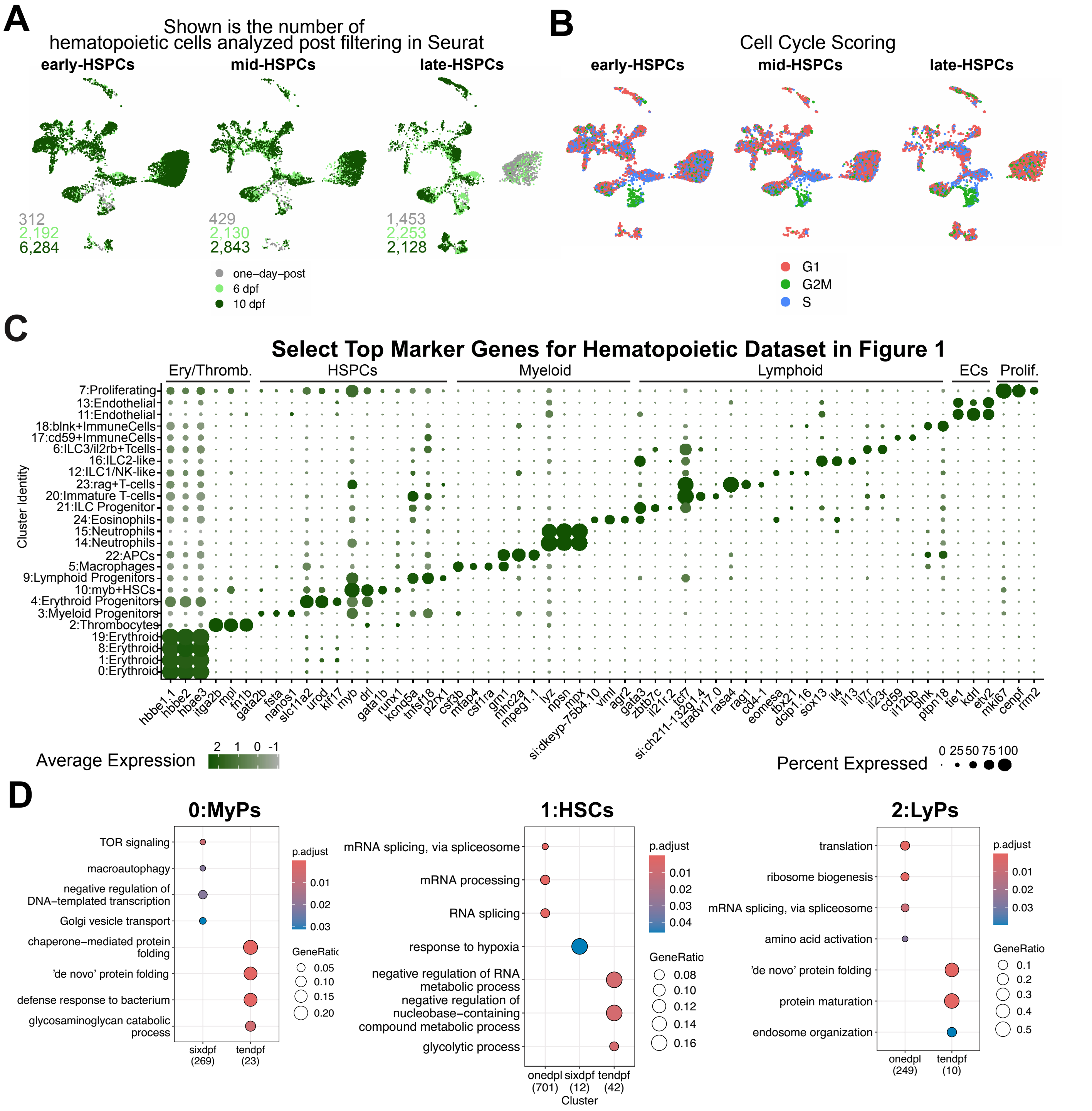


**Supplemental Figure 2. Characteristics of the hematopoietic populations.** (A) UMAP of 25 hematopoietic clusters split by trace and developmental timepoint illustrating no batch effects. (B) UMAP of cell cycle scores illustrating clustering is not cell cycle driven. The high G2M cluster is annotated proliferating. (C) Dot plot showing select marker genes of each cluster used to establish cluster identity. (D) EnrighGO Biological Processes pathway enrichment analysis on MyPs, HSCs and LyPs showing select categories across developmental time.

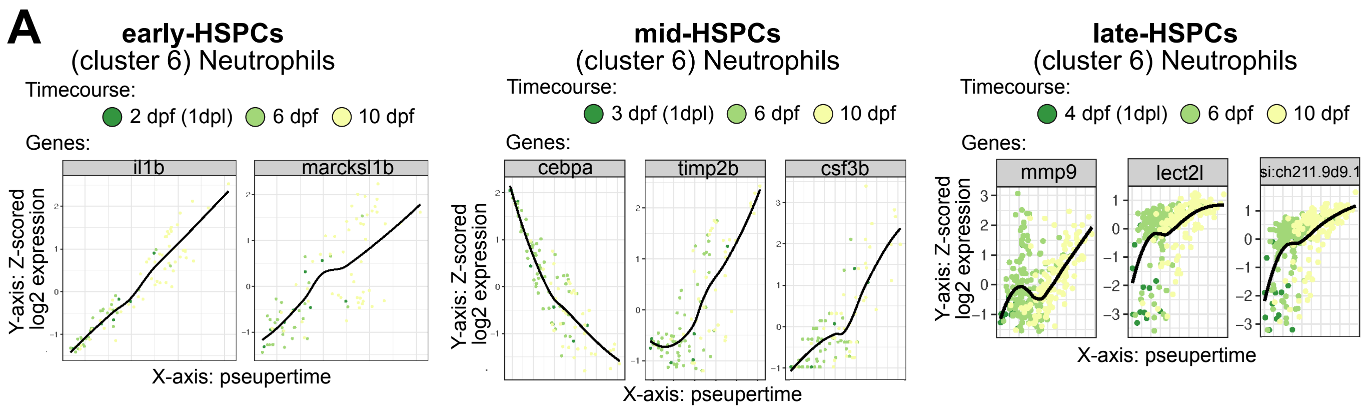


**Supplemental Figure 3. Psupertime analysis of neutrophils.** (A) Early- (left), mid- (middle), and late-HSPC (right) derived neutrophil cluster 14 along psupertime as shown in density plots with genes driving the alignment labeled above.

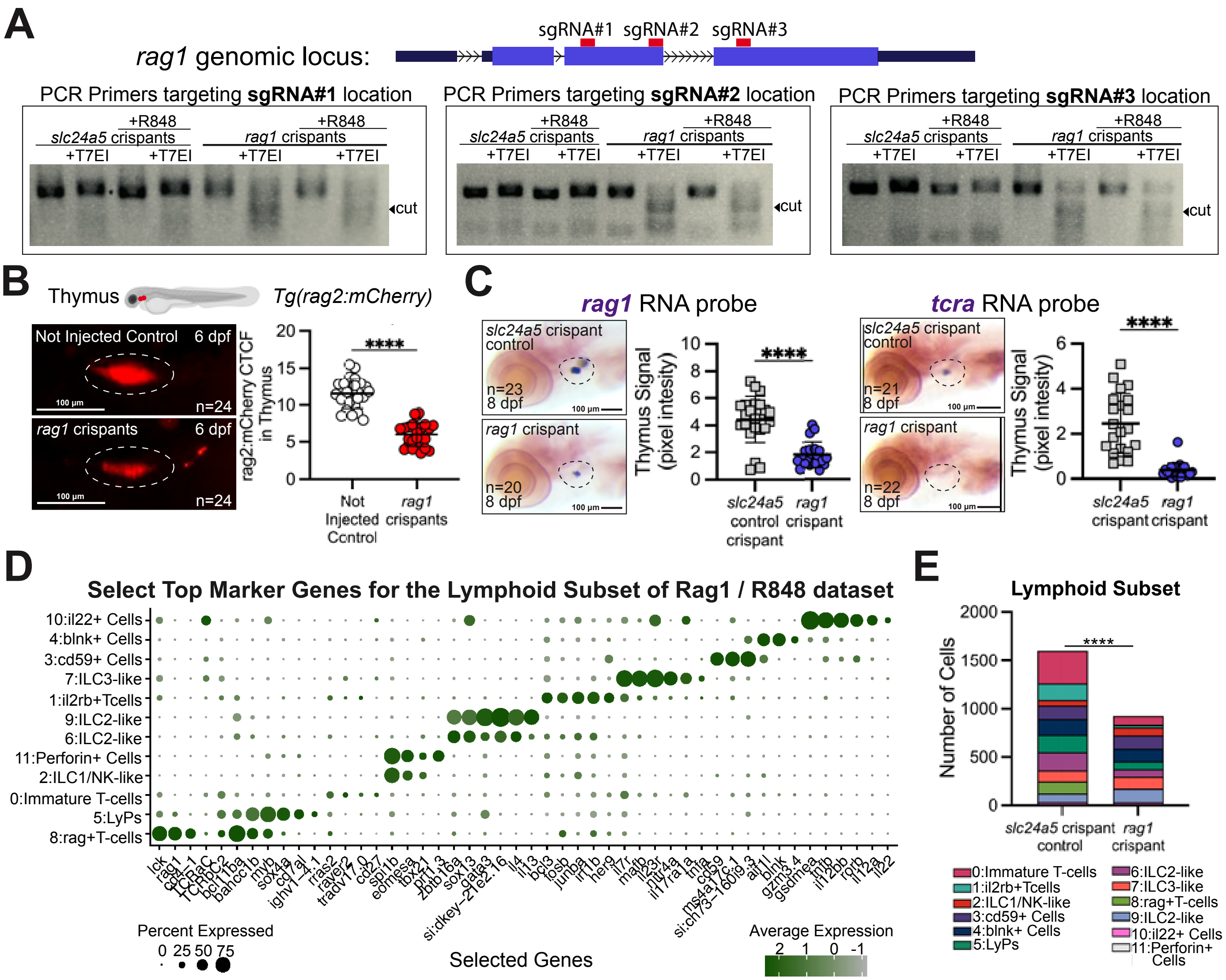


**Supplemental Figure 4. Quality control for *rag1* crispants.** (A) Schematic of the *rag1* locus, with exons in purple, and location of guide RNA targets in red (top). Gel images (bottom) of T7 endonuclease assay (T7EI) for mutagenesis showing effectiveness of *rag1* crispants approach. (B) Images of T-lymphocyte reporter *rag2:mCherry* (left) in the *rag1* crispants. Graph (right) depicting diminished signal in *rag1* crispants. Each point represents one fish. Significance computed with unpaired t-test. (C) RNA *In situ* hybridization showing significant reduction in *rag1* (left) and *tcra* (right) RNA expression in the thymus of that *rag1* crispants at 8 dpf. Each point represents one fish. Significance computed with unpaired t-test. (D) Dot plot showing the expression of selected genes in each cluster. (E) Bar plot showing the absolute number of cells per cluster shown in Main Figure 4. Significance computed with chi-square test. *-p<0.05, **p ≤0.01, **** p ≤0.0001

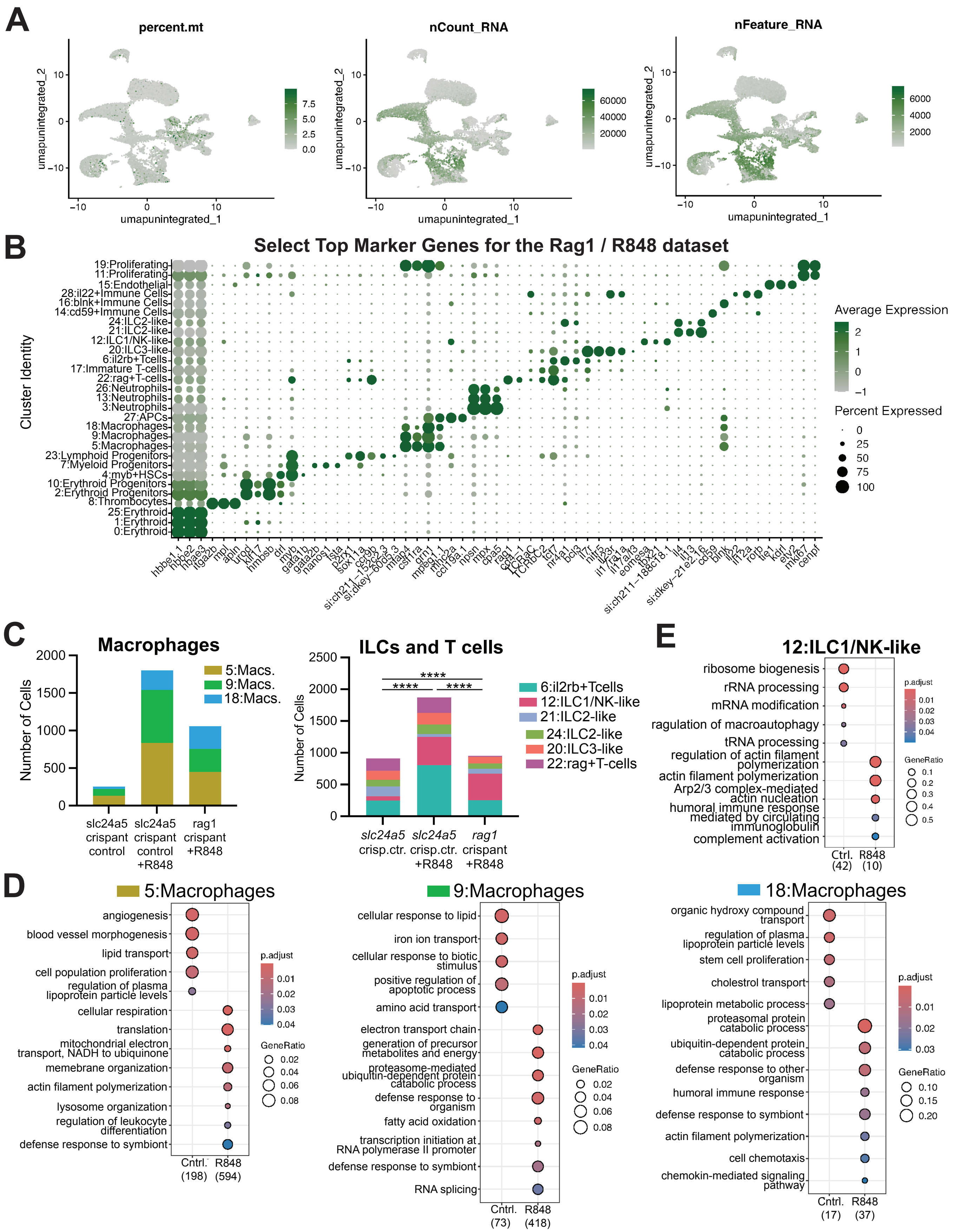
A
percent.mt
nCount_RNA
nFeature_RNA
umapunintegrated_1
umapunintegrated_2
B
Select Top Marker Genes for the Rag1 / R848 dataset
Cluster Identity
19:Proliferating
11:Proliferating
15:Endothelial
28:il22+Immune Cells
16:blnk+Immune Cells
14:cd59+Immune Cells
24:ILC2-like
21:ILC2-like
12:ILC1/NK-like
20:ILC3-like
6:il2rb+Tcells
17:Immature T-cells
22:rag+T-cells
26:Neutrophils
13:Neutrophils
3:Neutrophils
27:APCs
18:Macrophages
9:Macrophages
5:Macrophages
23:Lymphoid Progenitors
7:Myeloid Progenitors
4:myb+HSCs
10:Erythroid Progenitors
2:Erythroid Progenitors
8:Thrombocytes
25:Erythroid
1:Erythroid
0:Erythroid
Average Expression
Percent Expressed
C
Macrophages
Number of Cells
5:Macs.
9:Macs.
18:Macs.
slc24a5 crispant control
slc24a5 crispant control +R848
rag1 crispant +R848
ILCs and T cells
6:il2rb+Tcells
12:ILC1/NK-like
21:ILC2-like
24:ILC2-like
20:ILC3-like
22:rag+T-cells
slc24a5 crisp.ctr.
slc24a5 crisp.ctr. +R848
rag1 crispant +R848
E
12:ILC1/NK-like
ribosome biogenesis
rRNA processing
mRNA modification
ragulation of macroautophagy
tRNA processing
regulation of actin filament polymerization
actin filament polymerization
Arp2/3 complex-mediated actin nucleation
humoral immune response mediated by circulating immunoglobulin
complement activation
Ctrl. (42)
R848 (10)
p.adjust
GeneRatio
D
5:Macrophages
angiogenesis
blood vessel morphogenesis
lipid transport
cell population proliferation
regulation of plasma lipoprotein particle levels
cellular respiration
translation
mitochondrial electron transport, NADH to ubiquinone
memebrane organization
actin filament polymerization
lysosome organization
regulation of leukocyte differentiation
defense response to symbiont
Cntrl. (198)
R848 (594)
9:Macrophages
cellular response to lipid
iron ion transport
cellular response to biotic stimulus
positive regulation of apoptotic process
amino acid transport
electron transport chain
generation of precursor metabolites and energy
proteasome-mediated ubiquitin-dependent protein catabolic process
defense response to organism
fatty acid oxidation
transcription initiation at RNA polymerase II promoter
defense response to symbiont
RNA splicing
Cntrl. (73)
R848 (418)
18:Macrophages
organic hydroxy compound transport
regulation of plasma lipoprotein particle levels
stem cell proliferation
cholestrol transport
lipoprotein metabolic process
proteasomal protein catabolic process
ubiquitin-dependent protein catabolic process
defense response to other organism
humoral immune response
defense response to symbiont
actin filament polymerization
cell chemotaxis
chemokin-mediated signaling pathway
Cntrl. (17)
R848 (37)

**Supplemental Figure 5. Characteristics of the Rag1/R848 dataset.** (A) Feature plots showing percent mitochondria genes (left) ranging from 0-10%, number of counts (middle), and number of features or genes (right) expressed in each cell and cluster. (B) Dot plot showing select gene expression across the clusters in the Rag1/R848 dataset. (C) Bar plots illustrating the number of macrophages (left), ILCs/T-cells (right) in control, *rag1* crispant, and R848 treatment. Significance computed with chi-square test. (D,E) EnrichGO Biological Processes pathway enrichment analysis on differentially expressed genes in clusters 5, 9, 18, and 12 between control and R848 treated larvae.

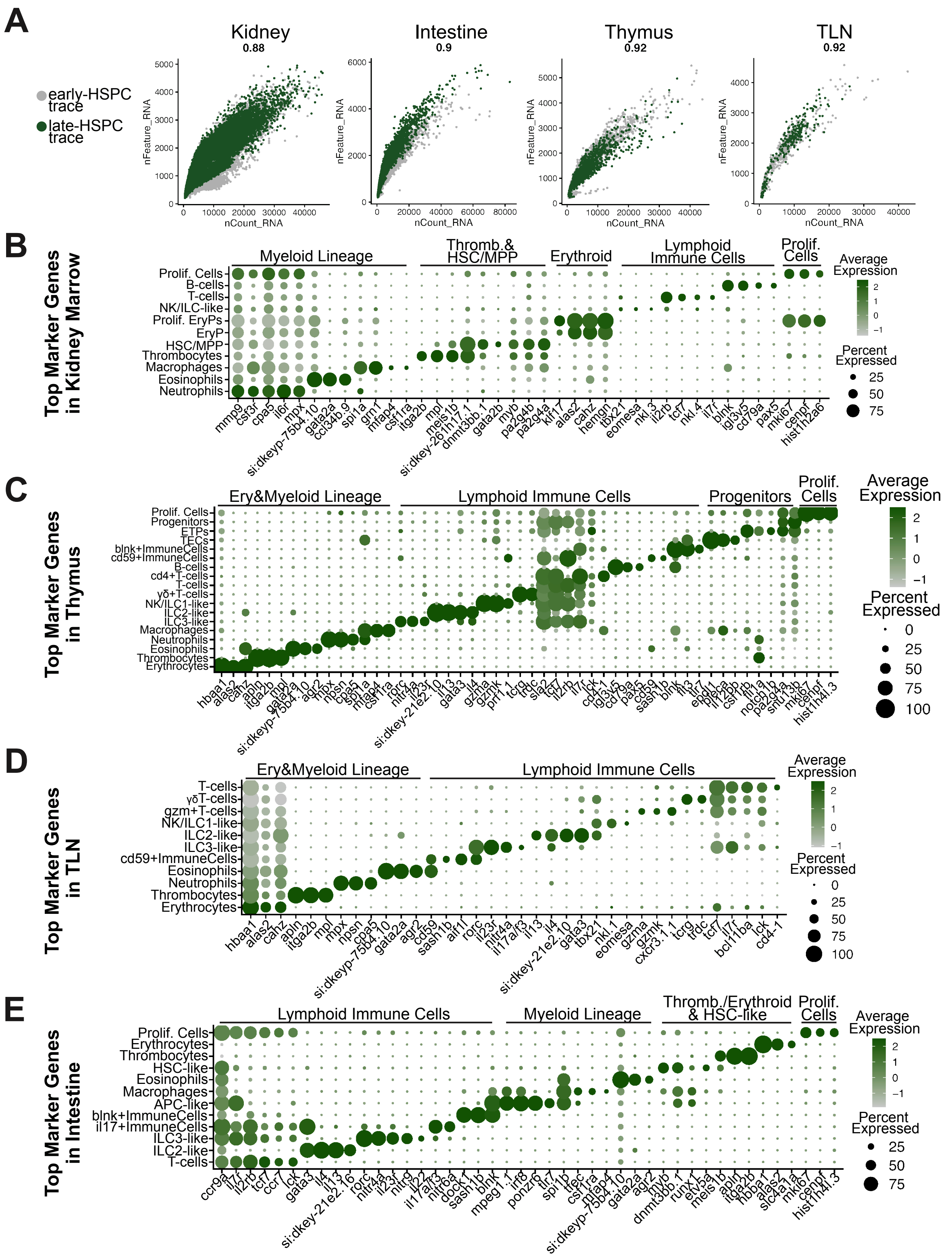
A
Kidney
0.88
Intestine
0.9
Thymus
0.92
TLN
0.92
early-HSPC trace
late-HSPC trace
nFeature_RNA
nCount_RNA
B
Top Marker Genes in Kidney Marrow
Myeloid Lineage
Thromb.& HSC/MPP
Erythroid
Lymphoid Immune Cells
Prolif. Cells
Prolif. Cells
B-cells
T-cells
NK/ILC-like
Prolif. EryPs
EryP
HSC/MPP
Thrombocytes
Macrophages
Eosinophils
Neutrophils
mmp9 csf3r cpa5 il6r mpx si:dkeyp-75b4.10 gata2a ccl34b.9 spi1a grn1 mfap4 csf1ra itga2b mpl meis1b si:dkey-261h17.1 dnmt3bb.1 gata2b myb pa2g4b pa2g4a klf17 alas2 cahz hemgn tbx21 eomesa nkl.3 il2rb tcf7 nkl.4 il7r blnk igl3v5 cd79a pax5 mki67 cenpf hist1h2a6
Average Expression
Percent Expressed
C
Top Marker Genes in Thymus
Ery&Myeloid Lineage
Lymphoid Immune Cells
Progenitors
Prolif. Cells
Prolif. Cells
Progenitors
ETPs
TECs
blnk+ImmuneCells
cd59+ImmuneCells
B-cells
cd4+T-cells
T-cells
γδ+T-cells
NK/ILC1-like
ILC2-like
ILC3-like
Macrophages
Neutrophils
Eosinophils
Thrombocytes
Erythrocytes
Average Expression
Percent Expressed
D
Top Marker Genes in TLN
Ery&Myeloid Lineage
Lymphoid Immune Cells
T-cells
γδT-cells
gzm+T-cells
NK/ILC1-like
ILC2-like
ILC3-like
cd59+ImmuneCells
Eosinophils
Neutrophils
Thrombocytes
Erythrocytes
hbaa1 alas2 cahz apln itga2b mpl mpx npsn cpa5 si:dkeyp-75b4.10 gata2a agr2 cd59 sash1b aif1l rorc il23r nitr4a il17a/f3 il13 il4 si:dkey-21e2.10 gata3 tbx21 nkl.1 eomesa gzma gzmk cxcr3.1 tcrg trdc tcf7 il7r bcl11ba lck cd4-1
Average Expression
Percent Expressed
E
Top Marker Genes in Intestine
Lymphoid Immune Cells
Myeloid Lineage
Thromb./Erythroid & HSC-like
Prolif. Cells
Prolif. Cells
Erythrocytes
Thrombocytes
HSC-like
Eosinophils
Macrophages
APC-like
blnk+ImmuneCells
il17+ImmuneCells
ILC3-like
ILC2-like
T-cells
ccr9a il7r il2rb tcf7 ccr7 lck gata3 il4 il13 si:dkey-21e2.16 rorc nitr4a il23r nitr9 il22 il17a/f3 nitr6a dock1 sash1b blnk mpeg1.1 irf8 ponzr6 tlr7 spi1b tfec csf1ra mfap4 si:dkeyp-75b4.10 gata2a agr2 myb dnmt3bb.1 runx1 etv5a meis1b apln itga2b hbba1 alas2 slc4a1a mki67 cenpf hist1h4l.3
Average Expression
Percent Expressed

**Supplemental Figure 6. Characteristics of the adult organ datasets.** (A) Scatter plots showing number of counts versus features by HSPC-trace for each organ. (B-E) Dot plots for each organ showing expression of select genes in each cluster. Genes were selected from top genes generated using FindAllMarkers function in Seurat with Wilcoxon Rank Sum testing.

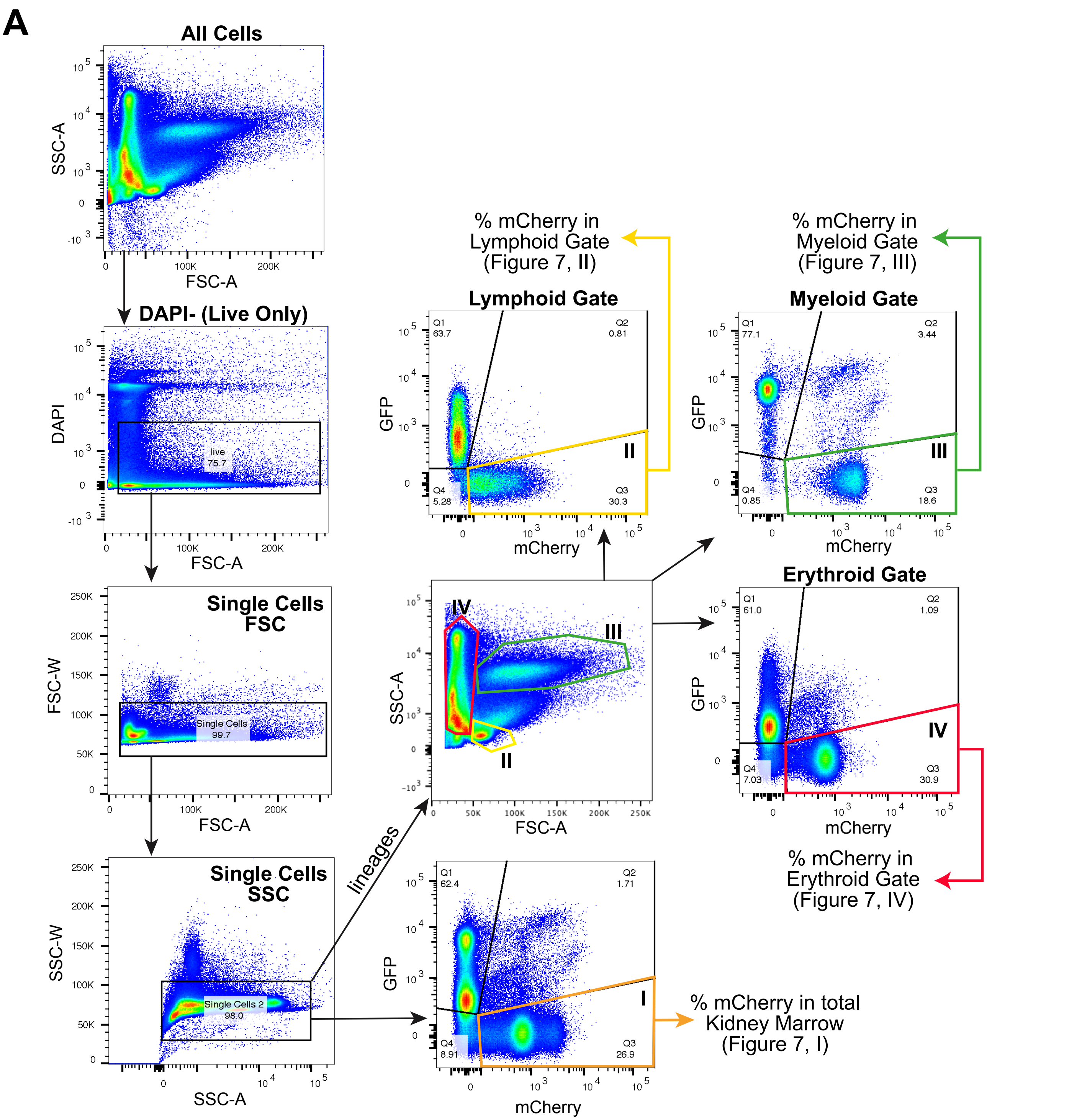


**Supplemental Figure 7. Gating strategy for the lineage tracing in *runx1* mutants.** (A) All cells were first filtered for live cells via DAPI dye exclusion. Singlets were selected by forward and side scatter. For quantification of mCherry+ cells in the total kidney marrow (orange gate- I), live single cells were analyzed by GFP versus mCherry to select for mCherry single positive cells representing fully switched cells. For defining early-, mid-, and late-HSPC trace contributions to each lineage, live, singlets were gated on lymphoid (II), myeloid (III) and erythroid (IV). Each lineage was analyzed by GFP versus mCherry to define the mCherry single positive percentage per lineage.

### SUPPLEMENTAL VIDEOS

**Supplemental Video 1:** Time lapse video of 6dpf larvae treated with vehicle control.

**Supplemental Video 2:** Time lapse video of 6dpf larvae treated with R848.

### SUPPLEMENTAL TABLES

**Supplemental Table 1.** Top genes and number of cells for all clusters in the early-, mid-, and late-HSPC trace scRNA-seq dataset.

**Supplemental Table 2.** Top genes, number of cells, DEGs, and GO analysis for the HSPC subset from the early-, mid-, and late-HSPC trace scRNA-seq dataset.

**Supplemental Table 3.** Top genes, cell numbers, DEGs, and GO analysis for the Myeloid-Lymphoid subset in the early-, mid-, and late-HSPC trace scRNA-seq dataset.

**Supplemental Table 4.** Top genes, cell numbers, DEGs, and module scores for the Lymphoid subset in the early-, mid-, and late-HSPC trace scRNA-seq dataset.

**Supplemental Table 5.** Top genes, cell numbers, and pathway analyses for the *rag1* crispant and R848 treatment scRNA-seq dataset.

**Supplemental Table 6.** Top genes and cell numbers for all clusters from the adult organs scRNA-seq datasets of early- and late-HSPC traced cells.

**Supplemental Table 7.** Pathway analyses and upstream motif analysis on the DEGs between early- and late-HSPC trace derived adult kidney marrow HSCs/MPPs.
